# Dynamics and variability in the pleiotropic effects of adaptation in laboratory budding yeast populations

**DOI:** 10.1101/2021.06.24.449852

**Authors:** Christopher W. Bakerlee, Angela M. Phillips, Alex N. Nguyen Ba, Michael M. Desai

**Affiliations:** Department of Molecular and Cellular Biology, Harvard University, Cambridge, MA, USA; Department of Organismic and Evolutionary Biology, Harvard University, Cambridge, MA, USA; Department of Cell and Systems Biology, University of Toronto, Toronto, Canada; Department of Physics, Harvard University, Cambridge, MA, USA; NSF-Simons Center for Mathematical and Statistical Analysis of Biology, Harvard University, Cambridge MA 02138; Quantitative Biology Initiative, Harvard University, Cambridge MA 02138

## Abstract

Evolutionary adaptation to a constant environment is driven by the accumulation of mutations which can have a range of unrealized pleiotropic effects in other environments. These pleiotropic consequences of adaptation can influence the emergence of specialists or generalists, and are critical for evolution in temporally or spatially fluctuating environments. While many experiments have examined the pleiotropic effects of adaptation at a snapshot in time, very few have observed the dynamics by which these effects emerge and evolve. Here, we propagated hundreds of diploid and haploid laboratory budding yeast populations in each of three environments, and then assayed their fitness in multiple environments over 1000 generations of evolution. We find that replicate populations evolved in the same condition share common patterns of pleiotropic effects across other environments, which emerge within the first several hundred generations of evolution. However, we also find dynamic and environment-specific variability within these trends: variability in pleiotropic effects tends to increase over time, with the extent of variability depending on the evolution environment. These results suggest shifting and overlapping contributions of chance and contingency to the pleiotropic effects of adaptation, which could influence evolutionary trajectories in complex environments that fluctuate across space and time.

## INTRODUCTION

As a population adapts to a given environment, it accumulates mutations that are beneficial in that environment, along with neutral and mildly deleterious ‘hitchhiker’ mutations. Because these mutations can also affect fitness in other environments, adaptation will tend to lead to pleiotropic fitness changes in other conditions. These pleiotropic consequences of adaptation need not be negative: evolution in one condition can lead to correlated fitness increases in similar environments as well as fitness declines in more dissimilar conditions. It is also natural to expect these consequences to vary over shorter or longer evolutionary timescales. For example, after a sufficiently long time adapting to a single condition, we might expect a population to increasingly specialize to that condition at the expense of its fitness elsewhere.

Numerous laboratory evolution experiments (Jerison et al. 2020; Ostrowski, Rozen, and Lenski 2005; Leiby and Marx 2014; Kinsler, Geiler-Samerotte, and Petrov 2020; Jasmin, Dillon, and Zeyl 2012; Novak et al. 2006; Meyer et al. 2010; V. S. Cooper and Lenski 2000; Bailey and Kassen 2012; Schick, Bailey, and Kassen 2015; Anderson et al. 2011; Li, Petrov, and Sherlock 2019; Dillon et al. 2016) as well as empirical studies of natural variation in diverse model systems (Geiler-Samerotte et al. 2020; Wang et al. 2015; M. C. Hall, Basten, and Willis 2006; Mackay and Huang 2018) have analyzed the pleiotropic consequences of adaptation. These studies have found examples of specialization, as well as cases of correlated adaptation and the evolution of more generalist phenotypes (Meyer et al. 2016; A. R. Hall, Scanlan, and Buckling 2011; Duffy, Turner, and Burch 2006; Duffy, Burch, and Turner 2007; Jerison et al. 2020; Li, Petrov, and Sherlock 2019; Leiby and Marx 2014). Pleiotropic fitness tradeoffs, such as those underlying specialization, can arise from either antagonistic pleiotropy (i.e., direct tradeoffs between the fitness effects of individual mutations across conditions), mutation accumulation (i.e., accumulation of mutations that are neutral in the evolution environment but impose fitness costs in other conditions), or some combination of these phenomena. More complex patterns of correlated fitness changes across conditions, such as those that underlie more generalist phenotypes, can result from more general relationships between fitness effects in different environments. Recent experimental and theoretical work has also analyzed how these distributions of mutational effects across environments can lead to an interplay between chance and contingency in determining both the typical pleiotropic consequences of adaptation and the predictability of these effects (Jerison et al. 2020; Ardell and Kryazhimskiy 2020).

The way in which these pleiotropic consequences of adaptation change as populations evolve is less well understood. That is, as a population adapts to a given environment, how steadily and consistently does its fitness change in alternate environments? Do these pleiotropic effects change systematically with time? For example, do fitness tradeoffs tend to become stronger the longer a population adapts to its home environment? And do the pleiotropic consequences of adaptation between replicate lines become more or less similar over time? These questions are critical both for understanding the nature of pleiotropic tradeoffs and for predicting the dynamics and outcomes of evolution in environments that fluctuate across time or space.

Previous studies have shed some light on these questions. For example, Meyer et al. (2010) reported on changes in phage susceptibility over 45,000 generations of *Escherichia coli* evolution, finding variable yet somewhat consistent trends across 6 evolved lines. Studying lines from the same evolution experiment, Leiby and Marx (2014) found a patchwork of pleiotropic patterns across 12 populations assayed for growth rate in 29 environments at two timepoints. While fitness changed predictably across replicates in some environments, changes were much more variable in others, with mutation rate modifying these patterns. However, these and other studies of the evolutionary dynamics of pleiotropy have been limited to a small number of timepoints, replicate populations, or evolution and assay environments (V. S. Cooper and Lenski 2000; Novak et al. 2006; Bailey and Kassen 2012). These limitations constrain the degree to which we can make useful inferences about how chance and contingency influence the pleiotropic consequences of adaptation, and how these consequences change over time.

To overcome these limitations, we experimentally evolved hundreds of uniquely barcoded haploid and diploid yeast populations in three environments for 1000 generations. Using sequencing-based bulk fitness assays, we assayed the fitness of each evolving population in five environments at 200-generation intervals spanning the 1000 generations of evolution. We then used the resulting data to quantify how the pleiotropic consequences of adaptation unfold in different evolution environments, along with the extent of variation among replicate populations. Our results allow us to investigate differential roles for chance and contingency over evolutionary time, with implications for the outcomes of adaptation in more complex fluctuating environments.

## RESULTS

To study the dynamics of the pleiotropic consequences of adaptation, we experimentally evolved 152 diploid yeast populations for about 1000 generations in one of three different environments (48 populations in YPD at 30°C, 54 populations in YPD + 0.2% acetic acid at 30°C, and 50 populations in YPD at 37°C). We chose these environments to facilitate comparisons with previous experimental evolution studies in yeast, which have used YPD at 30°C as a rich environment and acetic acid and high temperature to apply distinct types of stress (Nguyen Ba et al. 2019; Jerison et al. 2020). In addition, we evolved 20 haploid (MATα) yeast populations in YPD at 37°C; these are a subset of populations that did not autodiploidize from a larger haploid evolution experiment (see Methods for details).

Each haploid population was founded by a single clone of a putatively isogenic laboratory strain, labeled with a unique DNA barcode at a neutral locus prior to the evolution experiment (Fig. 1A). Diploid populations were founded by mating uniquely barcoded haploids and selecting for diploids. We then propagated each population for 1000 generations in batch culture, with a 1:2^10^ dilution every 24 hours; this corresponds to an effective population size of ~2 × 10^5^ (Fig. 1A; see Methods for details). We froze an aliquot from each population at 50-generation intervals at −80°C in 8% glycerol for long-term storage.

**Figure 1.**
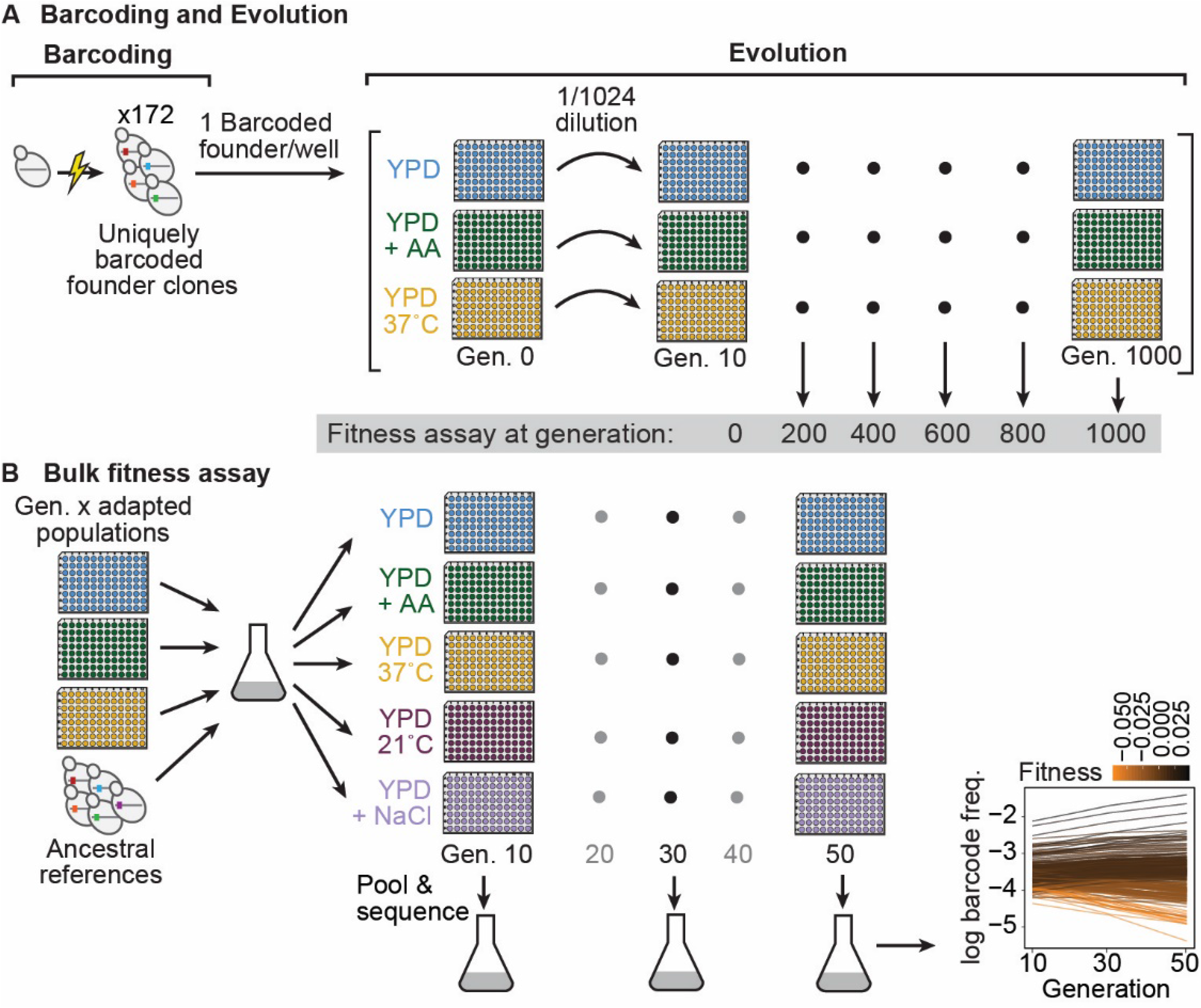
Evolution experiment and bulk fitness assay. **(A)** Yeast cells were uniquely barcoded to generate founder clones. Uniquely barcoded founder clones were used to seed individual populations in 96-well plates. Populations were evolved for 1,000 generations in three distinct environments: rich media (YPD), rich media at elevated temperature (YPD, 37°C), and rich media with 0.2% acetic acid (YPD + AA), and frozen at 50-generation intervals. Fitness assays were performed at 200-generation intervals. **(B)** Bulk fitness assay of barcoded adapted populations by competitive growth in each evolution environment and two additional environments (YPD, 21°C and YPD + 0.4 M NaCl). Relative fitness of each population was evaluated from the log frequency of the respective barcode sequence over time compared to that of ancestral references, based on assay generations 10, 30, and 50. ***Figure 1–figure supplement 1.*** Bulk fitness assay technical replicate fitness correlations.

After completing the evolution, we revived populations from generation 0, 200, 400, 600, 800, and 1000. We then conducted parallel bulk fitness assays (2 technical replicates) to measure the fitness of each population at each timepoint across five environments (the three evolution environments plus YPD + 0.4M NaCl at 30°C (transfers every 24 hours) and YPD at 21°C (transfers every 48 hours), environments which exposed the populations to unique osmotic and temperature stresses). In each bulk fitness assay, we pooled all populations from a given generation along with a small number of common reference clones and propagated them for 50 generations (Fig 1B). We then sequenced the barcode locus at generation 10, 30, and 50, and we inferred the fitness of each population from the change in log frequency of each corresponding barcode. By exploiting the fact that each population is uniquely barcoded, these bulk fitness assays allowed us to estimate the fitness of all 172 populations at each of the five 200-generation intervals in each of the five environments with minimal cost and effort (see Methods for details).

Based on the measured fitness of the generation 0 ancestral populations, we found that some diploid populations had substantially higher ancestral fitness in certain assay environments, likely because they acquired mutations prior to the start of the evolution. To clarify our downstream analyses, we excluded 19 outlier diploid populations whose ancestors differed from the mean ancestral fitness by at least 4% in at least one environment, leaving us with 133 diploid populations (43 YPD at 30°C, 48 YPD + acetic acid, and 42 YPD at 37°C) and 20 haploid populations (153 populations total). However, we note that the results of all our analyses are very similar when we consider the entire dataset with outliers included (see Figure Supplements).

### Adaptation to the home environment leads to consistent fitness gains and pleiotropic effects

While there is modest variability between replicate populations, adaptation in each environment leads to a consistent increase in fitness in that “home” environment (Fig. 2, subplots with bold black borders). As observed in earlier experiments, this fitness increase is largely predictable, and follows a characteristic pattern of declining adaptability: early rapid fitness gains that slow down over time (Couce and Tenaillon 2015). This declining adaptability trend is less obvious among populations evolved at 37°C, possibly because the fitness gains in this environment were generally minimal, but we do observe declining adaptability in the handful of diploid populations at 37°C that experienced larger-than-average fitness gains.

**Figure 2.**
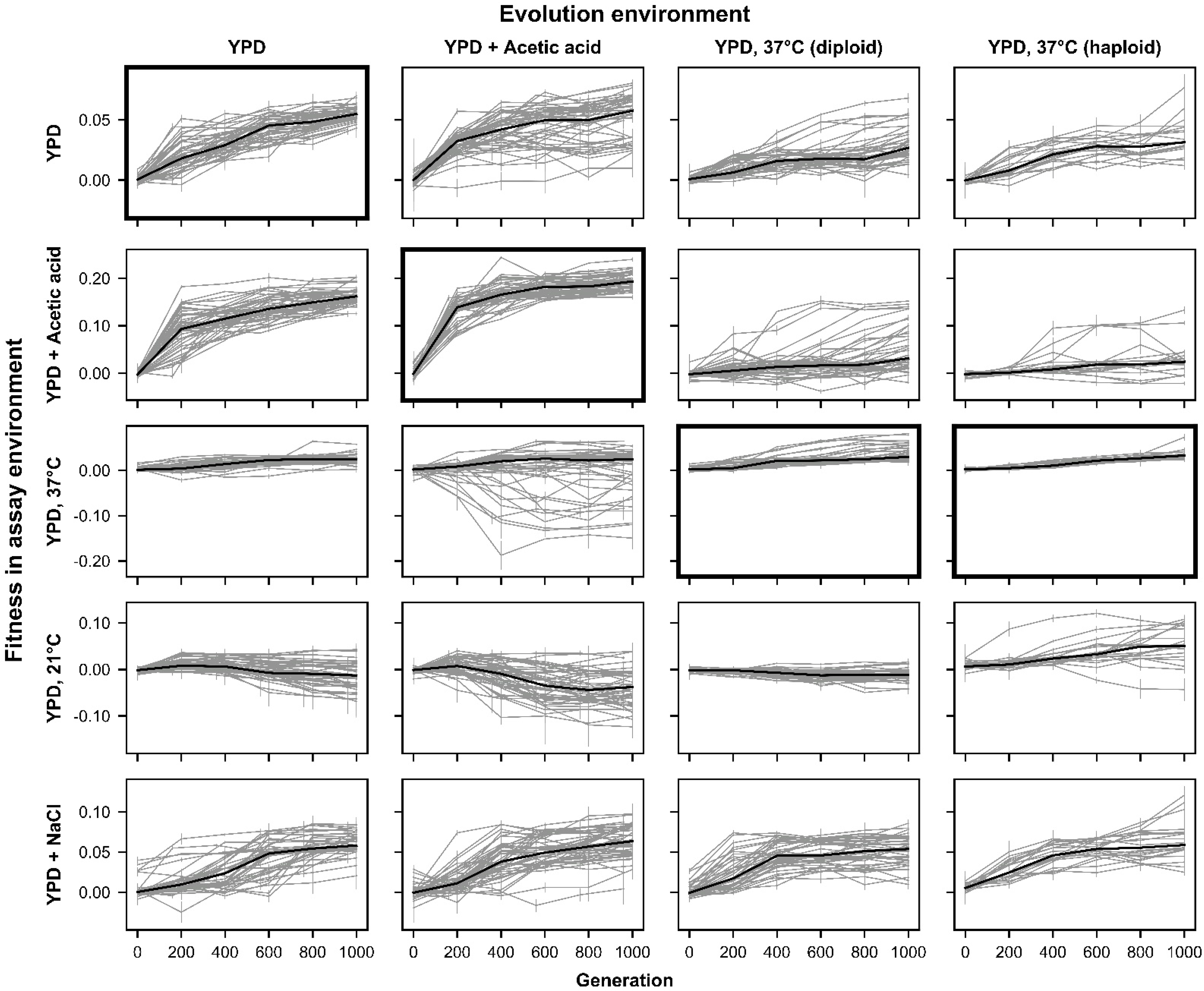
Fitness changes over 1000 generations of evolution. Replicate populations for each evolution condition are shown in each column. Environments in which these populations’ fitnesses were assayed are shown in the rows. Plots for which evolution and assay environment are the same are indicated by a bold outer border. The black line in each plot indicates the median fitness. Error bars indicate standard error of the mean. ***Figure 2–figure supplement 1.*** Fitness changes over 1000 generations of evolution for unfiltered data. ***Figure 2 – source data 1.*** Bulk fitness assay read counts and measured fitnesses.

Adaptation in each evolution environment also led to fitness changes in most other environments (Fig. 2). In general, these fitness changes tend to have a consistent direction over time for each environment pair. For example, populations adapted to YPD + acetic acid and YPD at 37°C steadily gained fitness in the YPD at 30°C and YPD + 0.4M NaCl environments over time, with the average fitness across populations largely following the same trend seen at home: initial rapid fitness gains followed by slower increases over time. In other instances, fitness gains at home correspond to fitness declines in away environments. For example, populations evolved in YPD + acetic acid tend to lose fitness in YPD at 21°C. However, pleiotropic effects are less predictable than the fitness gains in the home environment: we see more variability among replicate lines in away environments, both in the shapes of their fitness trajectories and in their ultimate evolutionary outcomes (e.g. some populations evolved in YPD + acetic acid in fact gain fitness in YPD at 21°C) (see analysis below).

To visualize how these pleiotropic effects change over time, we plot these fitness trajectories across pairs of environments (Fig. 3). This representation of the data shows clear but sometimes subtle differences in patterns of pleiotropy depending on evolution environment and ploidy. For instance, while almost all populations gained fitness in both YPD at 30°C and YPD + NaCl, the dynamics of fitness change differed based on evolution environment: populations evolved at 37°C (orange lines in Fig. 3) initially made substantial fitness gains in YPD + NaCl sometimes followed by more significant gains in YPD at 30°C, whereas the populations evolved in YPD at 30°C (cyan lines) and YPD + acetic acid (green lines) only gained substantial fitness in YPD + NaCl after initial fitness increases in YPD at 30°C (Supplementary File 1). Separately, plotting fitness in YPD + acetic acid against fitness in YPD at 21°C reveals trajectories that segregate not only by evolution environment, but also by ploidy (Supplementary File 1).

**Figure 3.**
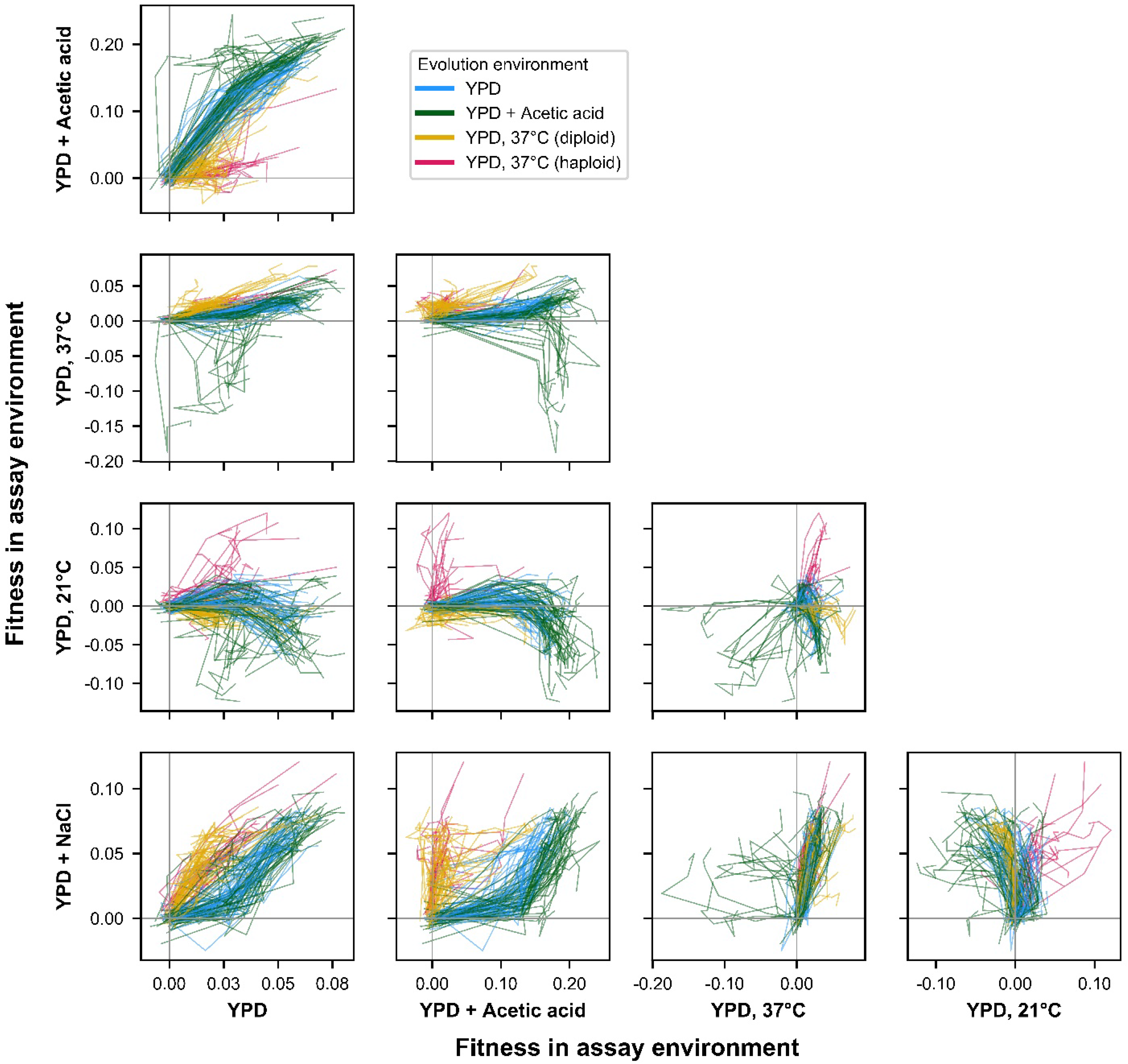
ExE evolutionary trajectories over 1000 generations of evolution in a constant environment. Axes correspond to fitness in the indicated assay environments. Colors correspond to evolution condition. Grey vertical and horizontal lines indicate zero fitness relative to an ancestral reference in each environment. ***Figure 3–figure supplement 1.*** ExE evolutionary trajectories over 1000 generations of evolution in a constant environment for unfiltered data.

### Characteristic environment- and ploidy-specific pleiotropic profiles emerge over time

To understand the diversity of fitness trajectories across environments, we treated the fitness of each population across all five assay environments as a single “pleiotropic profile.” We then conducted principal component analysis across all these pleiotropic profiles to characterize variation between replicate populations, across different evolution environments, and over time.

In Fig. 4A, we plot the first two principal components of each pleiotropic profile (which together consistently explain well over half the variance in the data (Fig. 4 -- figure supplement 2)) for populations from each of the six measured timepoints. We see that the populations separate over time into somewhat distinct clusters based on their evolution environment and ploidy. These clusters suggest that evolution in each environment leads to the formation of a characteristic environment- and ploidy-specific pleiotropic profile.

**Figure 4.**
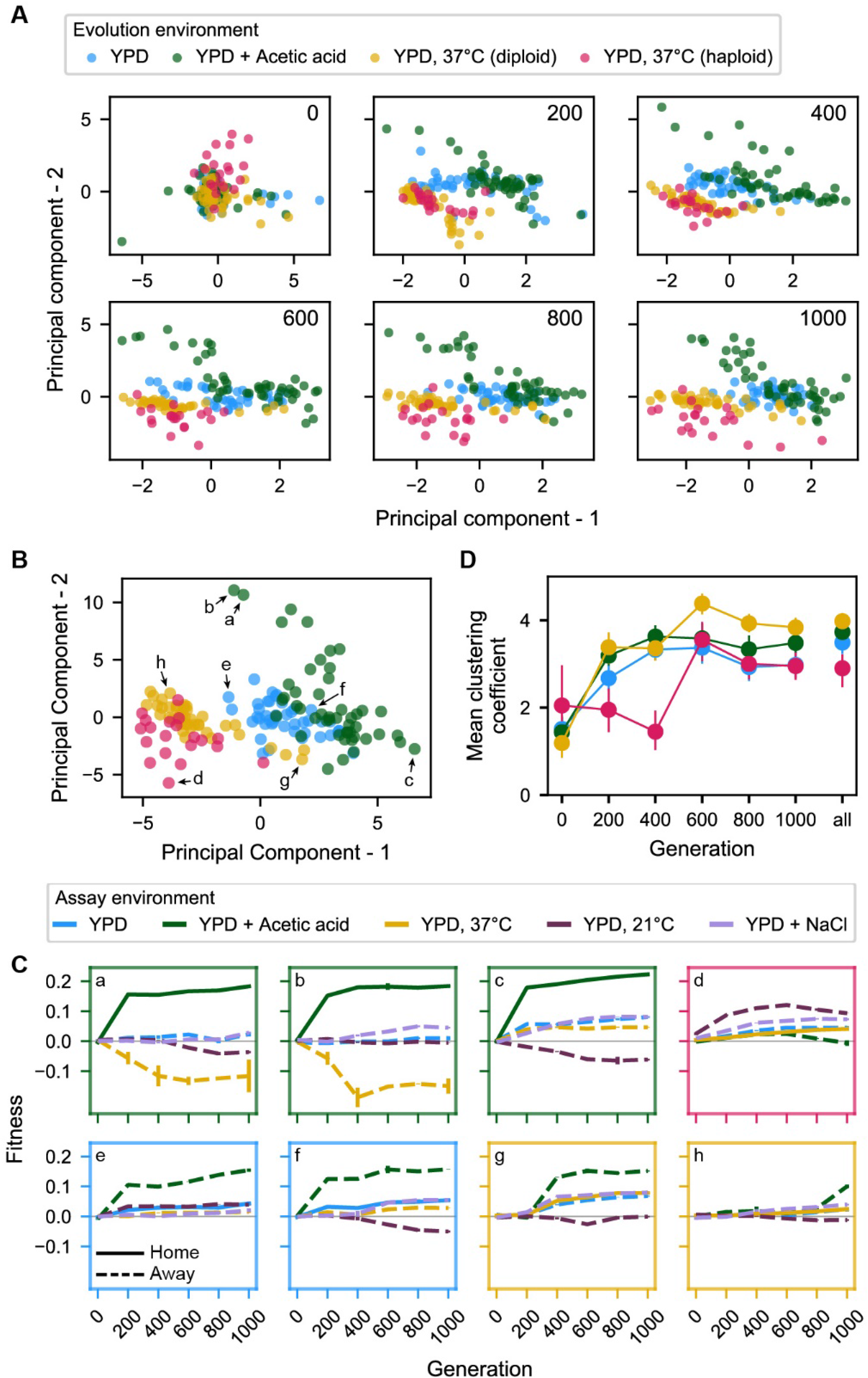
Principal component analysis of pleiotropy. **(A)** Principal component analysis of evolving populations, performed independently each 200 generations. The first two PCs are plotted. Populations are colored according to evolution condition. **(B)** Principal component analysis of all populations using all fitness data from across the 1000 generations. The first two PCs are plotted and explain 30% and 22% of the variance, respectively. **(C)** Plots of fitness trajectories in all 5 assay environments for 8 example populations (a-h, identified as points in (B)). **(D)** Population clustering in PCA by evolution condition over time. Clustering of each population was quantified as the number of five nearest neighbors that share the same evolution condition, for each 200-generation interval, and across all intervals. Clustering metrics were averaged for each evolution condition to calculate point estimates; error bars represent 95% confidence intervals of the mean clustering metric, estimated by performing PCA on bootstrapped replicate fitness measurements. ***Figure 4–figure supplement 1.*** Principal component analysis of pleiotropy for unfiltered data. ***Figure 4 – figure supplement 2.*** Variance explained by principal components in (A) and the corresponding panel of figure supplement 1. ***Figure 4 – figure supplement 3.*** Relative contributions of each interval to principal components in (B) and the corresponding panel of figure supplement 1. ***Figure 4 – figure supplement 4.*** Population clustering in PCA as in (D) quantified for three and ten nearest neighbors. ***Figure 4 – source data 1.*** Principal component analyses presented in Figure 4A. ***Figure 4 – source data 2.*** Principal component analysis presented in Figure 4B. ***Figure 4 – source data 3.*** Principal component analyses presented in Figure 4 – figure supplement 1A. ***Figure 4 – source data 4.*** Principal component analysis presented in Figure 4 – figure supplement 1B.

Characteristic pleiotropic profiles can also be observed when running principal component analysis on the complete concatenated (but unordered) fitness data (i.e., with the pleiotropic profile of each population now defined as its fitness across all five assay environments at all six 200-generation timepoints, a total of 30 measurements) and plotting data according to the first two components, which explain 30% and 22% of total variance, respectively (Fig. 4B). To provide an intuition for the meaning of distance and location in this principal component space, we show home and away environment fitness trajectories for select populations indicated in Figure 4B (Fig. 4C). The extent of evolution condition-specific clustering in this two-dimensional PCA is indicative of characteristic pleiotropic profiles (Fig. 4C), and it appears comparable to that observed in analyses conducted independently for generations 600, 800, and 1000. This is unsurprising given the outsized weighting of later generations in each principal component (Fig. 4 -- figure supplement 3).

To more formally quantify the emergence of characteristic pleiotropic profiles over time in Figures 4A and B, we developed a simple clustering metric, which counts how many of a given population’s five nearest neighbors belong to the same evolution condition on average. We see that the degree of clustering in this two-dimensional space rises appreciably until the 600-generation mark, at which point it plateaus (Fig. 4D). The observed clustering from generation 200 onward is much greater than expected by chance, as is clustering for the total-data PCA shown in Figure 4B (compared to a null expectation constructed by randomly permuting the evolution condition assigned to each population; *p* < 0.001). Note that this trend is consistent when the number of neighbors in the analysis is lowered to 3 or elevated to 10 (Figure 4—figure supplement 4). Thus, we observe the rapid emergence and later stabilization of general pleiotropic profiles characteristic to each evolution condition.

### General trends contain significant variation, which varies with ploidy, environment, and time

Our principal component analysis shows that replicate populations in each evolution condition tend to follow similar trends in fitness changes across environments, leading to characteristic environment-specific pleiotropic profiles. However, it is apparent from Fig. 2 and Fig. 3 that there remains significant stochastic variability in the pleiotropic effects of adaptation among populations evolved in the same environment. For instance, populations evolved in the acetic acid environment splay out into all four quadrants when plotting fitness at 37°C against fitness at 21°C (Fig 3; Supplementary File 1). This variability can also be seen in the wide dispersion of populations within clusters in Fig. 4B, particularly among diploids evolved in the acetic acid environment and at 37°C.

We find that these patterns of variability are structured, with specific evolution conditions fostering more variable outcomes in certain assay environments (Fig. 5). For example, populations evolved in YPD + acetic acid exhibit generally wider variation in home and away environments than populations evolved in other environments. While it is tempting to link this pattern to the large fitness gains these populations make in their home environment, we note that populations evolved in YPD at 30°C also make significant correlated gains in YPD + acetic acid without generating such variable results across other assay environments. This suggests that, with respect to the distribution of pleiotropic effects of fixed driver or hitchhiking mutations, paths to higher fitness in YPD + acetic acid are qualitatively different for the populations evolved in YPD at 30°C. In another example, while diploid and haploid populations evolved at 37°C show similar variability in 37°C, 30°C, and YPD + NaCl across the experiment, they experience more variable outcomes in YPD + acetic acid and 21°C, respectively. Together, these results suggest that the role for chance in the pleiotropic trajectories of evolving populations is contingent on the condition to which the population is adapted.

**Figure 5.**
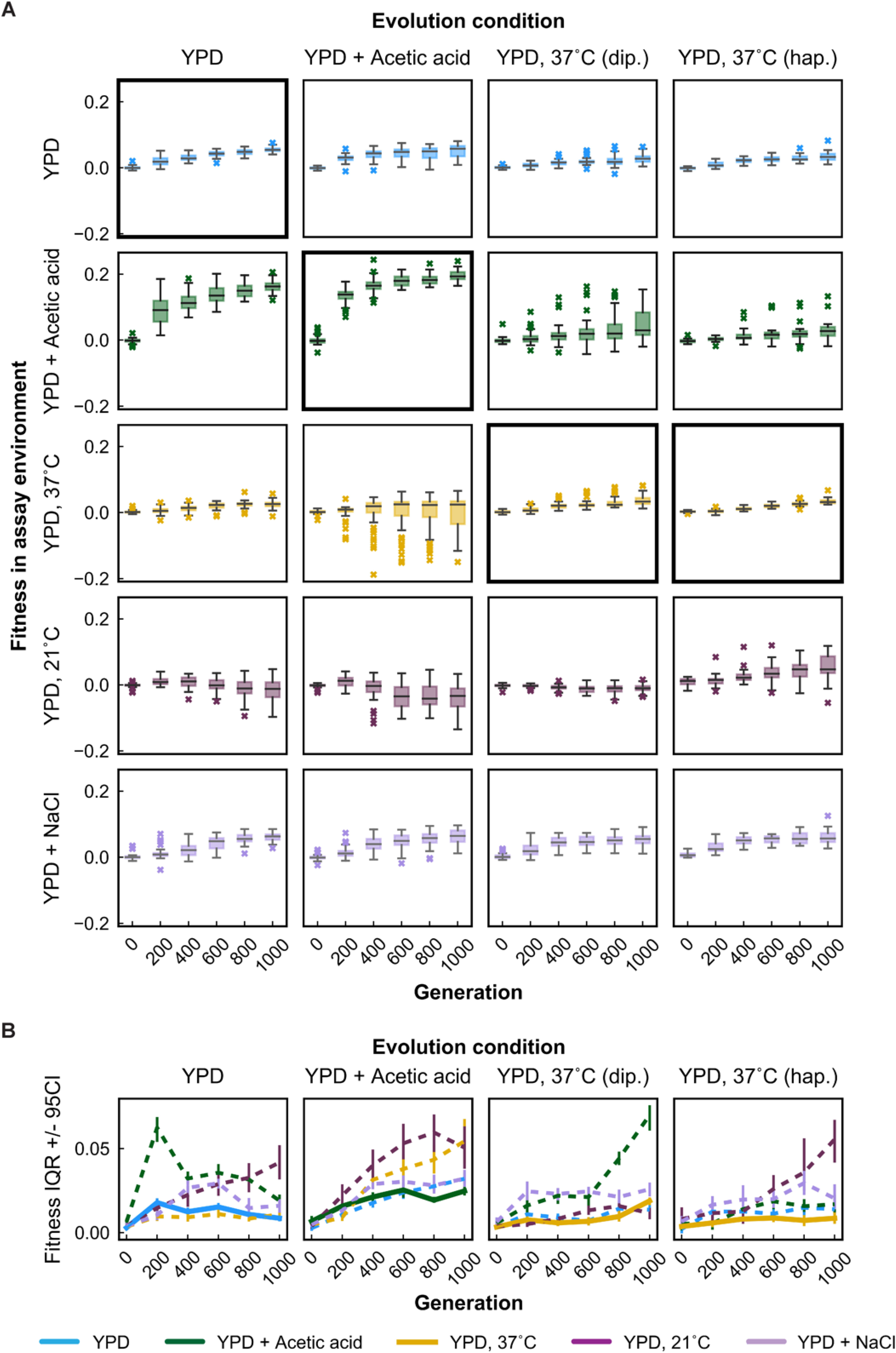
Variability in fitness over time. **(A)** Box plots summarizing population mean fitness over time for each evolution condition (columns) in each assay environment (rows). Line, box, and whiskers represent the median, quartiles, and data within 1.5xIQR of each quartile, respectively; outlier populations beyond whiskers are shown as points. **(B)** IQR from box plots in (A) are plotted as a function of time for each evolution condition and assay environment. IQR for fitness measured in home and away environments are represented by solid and dashed lines, respectively. Error bars represent 95% confidence intervals of IQR calculated from bootstrapped replicate fitness measurements. ***Figure 5–figure supplement 1***. Variability in fitness over time for unfiltered data. ***Figure 5–figure supplement 2***. Brown-Forsythe significance test results for differences between variance at home and away.

In addition, the variation in outcomes is a function of evolutionary time. While variation in fitness at home tends to remain relatively low over the course of 1000 generations (Fig. 5A, bold black boxes; Fig 5B, thick solid lines), variation in away environments generally (if haltingly) increases over time, with a few exceptions. In other words, selection appears to suppress variation among trajectories in the home environment, at least on the timescales studied. To assess the statistical significance of these differences in variance, we used a one-tailed variant of a Brown-Forsythe test to perform pairwise comparisons of home and away fitness variance among replicate lines evolved in a given condition at each evolution timepoint. Of the 80 non-ancestral pairwise comparisons, over half (48) indicated significantly greater variance in the away environment (at a threshold of *p* < 0.05) and only 6 showed significantly greater variance at home (Figure 5—figure supplement 2).

The role of stochasticity and temporal shifts in pleiotropic dynamics also can be seen in the relative non-monotonicity of fitness trajectories in away environments compared to home environments. To assess non-monotonicity, we interpolated fitness at 500 generations for each population in each assay environment and compared the 0-to-500-generation and 500-to-1000-generation fitness changes. Trajectories were considered non-monotonic if fitness changes in these intervals were in opposite directions (Fig. 6A, see shaded quadrants), reflecting pleiotropic effects that change in sign over time. We find that populations rarely possess clearly non-monotonic trajectories in their home environment (4/153 trajectories, or 2.6%), whereas they much more commonly (*p* < 0.0001, χ2 test) possess clearly non-monotonic trajectories in away environments (102/612 trajectories, or 16.7%) (Fig. 6B). Many but not all of these monotonic trajectories (72/102, or 71%) reflect initially positive pleiotropic effects that become negative in the second half of the experiment, as we might expect if a population increasingly specializes to its home environment over time.

**Figure 6.**
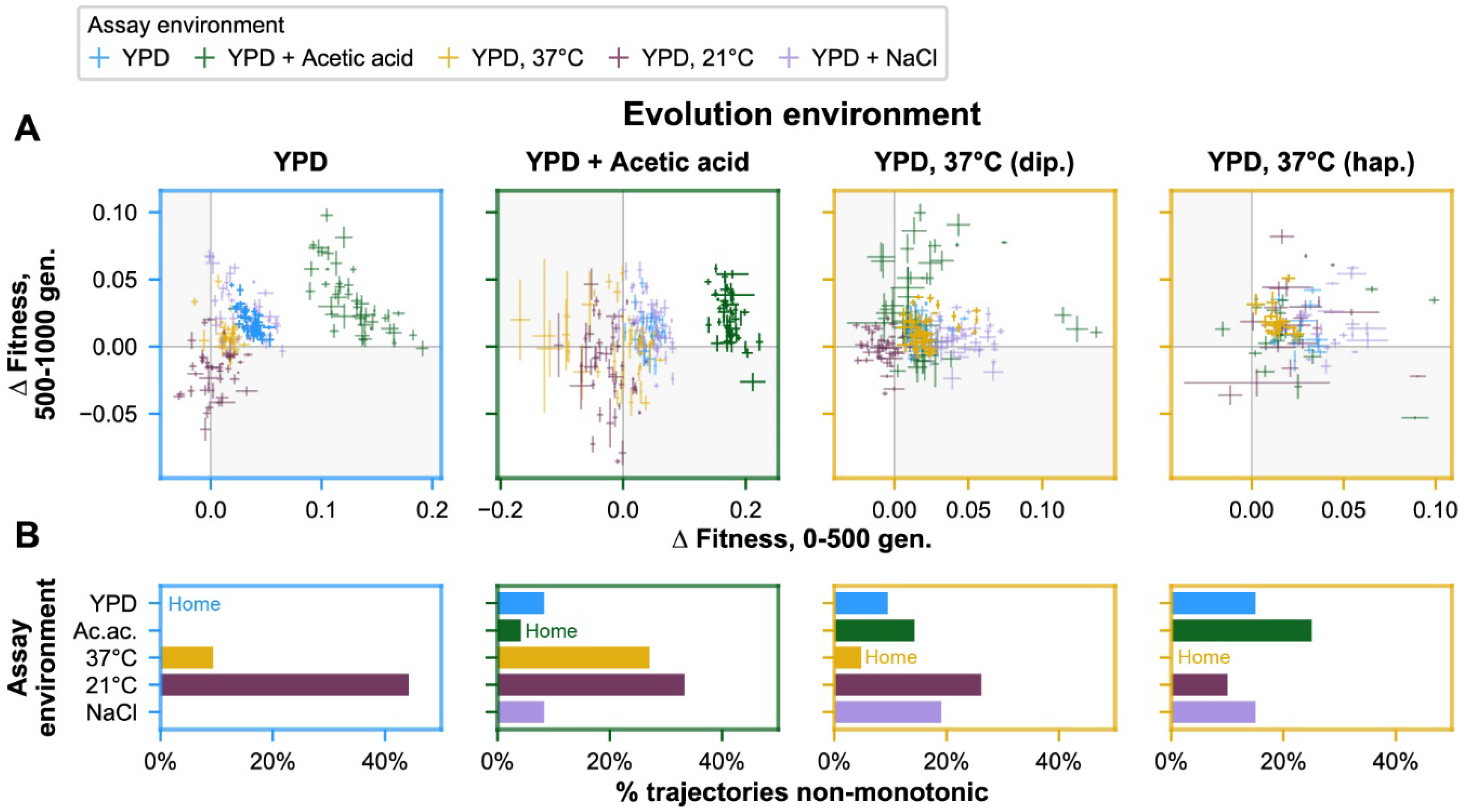
Non-monotonicity in evolutionary trajectories. **(A)** Each panel shows, for each of the 5 assay environments, the change in fitness over the first 500 (x-axis) and second 500 (y-axis) generations of evolution of each population in a given evolution environment. Populations that fall in shaded quadrants have trajectories that are non-monotonic. Points corresponding to fitness in the home environment are colored more opaquely than points corresponding to fitness in away environments, and panel borders have been colored to match the home environment. Fitness at generation 500 has been interpolated. **(B)** Each panel corresponds to a given evolution environment and shows the proportion of populations evolved in that environment that exhibit clearly non-monotonic fitness trajectories in (A). “Clearly non-monotonic” trajectories are those populations (points) in (A) that fall in the grey quadrants and whose error bars (1 standard error in either direction) do *not* span either the x- or y-axis. As in (A), bars corresponding to the home environment are colored more opaquely than bars corresponding to away environments. ***Figure 6–figure supplement 1.*** Non-monotonicity in evolutionary trajectories for unfiltered data.

## DISCUSSION

To characterize the dynamics of pleiotropy during adaptation, we evolved hundreds of diploid and haploid yeast populations in three environments for 1000 generations, and assayed their fitness in these and two other environments at 200-generation intervals. Our results offer insight into how pleiotropic effects emerge and change on an evolutionary timescale. Consistent with earlier work, we observe repeatable fitness trajectories across many replicate populations in their home environments, which follow a pattern of initial rapid fitness gains followed by declining adaptability over time. Replicate populations also tend to follow consistent fitness trajectories in away environments, whether gaining or losing fitness on average. Looking across populations and environments, characteristic patterns of pleiotropy specific to each evolution condition emerge rapidly and stabilize within about 600 generations.

Despite these characteristic patterns, we also observe ample variability within these trends. Examining the fitness trajectories of populations individually, we find that about 17% of away-environment trajectories are non-monotonic, compared to just 3% of home-environment trajectories. This non-monotonicity is indicative of the sequential establishment of mutations with opposing pleiotropic effects in these populations. Meanwhile, across replicate populations, there is substantial variability in the pleiotropic consequences of evolution in each condition. Consistent with past work, we observe more variability in away than in home environments at the end of the experiment (Travisano and Lenski 1996; Ostrowski, Rozen, and Lenski 2005). However, our results also reveal how populations can follow very different trajectories in arriving at these endpoint fitnesses. Diverse away-environment trajectories manifest as changes in the variance among replicate populations over time, with a general tendency for variance to increase over the course of the experiment.

Together, patterns of pleiotropy along with variability among replicate populations suggest an important and dynamic role for chance and contingency in the fates of populations evolving in environments that fluctuate in space and time. Whether populations trend toward specialist or generalist phenotypes will not simply reflect physiological constraints (Bono et al. 2017; Jerison et al. 2020). Rather, as we observe, mutational opportunities to move toward higher or lower fitness in alternate environments may be accessible at all times. Thus, the emergence of specialism or generalism will be a product of both the distribution of pleiotropic effects of mutations that establish and dynamical factors that influence the timescale, sequence, and likelihood of their fixation (e.g., epistasis, ploidy, clonal interference, mutation rate, population size).

Furthermore, the timescale over which pleiotropic effects emerge and change will interact with patterns of environmental fluctuations to determine evolutionary outcomes. In the conditions studied here, we observe that pleiotropic profiles generally emerge early and stabilize by 600 generations. Independent of other dynamical consequences of the rate of environmental change (Cvijović et al. 2015), it is therefore likely that fluctuations on longer timescales (e.g., longer than 600 generations in this system) will lead to qualitatively different outcomes than fluctuations on shorter timescales. Our data show that both the average and variance in these outcomes will also depend critically on the specific sequence of environments experienced by a population.

These results underscore the need for further empirical and theoretical work to understand patterns of pleiotropic effects over time and their effects on evolutionary trajectories. Additional experiments will be required to describe how general pleiotropic trends and variability within these trends arise and shift across a wider array of environments, as well as in different model systems. Likewise, studies of pleiotropy in populations evolved for longer periods, such as those described by Johnson et al. (2021), may provide a richer perspective on the repeatability, diversity, and stability of pleiotropic trajectories. Finally, this work motivates further theoretical inquiry into how the dynamics and variability of pleiotropic effects will interact with other important parameters -- such as patterns of environmental fluctuation, mutation rate, sexual recombination, and the underlying distributions of fitness effects -- to influence evolutionary outcomes. Integrating empirical datasets like the one presented here with such theoretical insight will enable better prediction of adaptation in complex environments.

## MATERIALS AND METHODS

### Strain generation

Strains in this study are derived from YAN404 and YAN407 (Nguyen Ba et al. 2019), which were constructed on the BY4742 background (S288C: *MATα*, *his*3Δ1, *ura*3Δ0, *leu*2Δ0, *lys*2Δ0) to add the *RME1*pr::ins-308A mutation, meant to improve transformation efficiency in both the *MAT****a*** and *MAT****α*** cell types. Several additional modifications were made to enable proper barcoding, mating, and selection, as stated in Supplementary File 2. Ultimately, YCB140B and YCB137A (and YCB140B × YCB137A mated diploids) were used to found the populations evolved in this experiment.

### Barcode plasmid design and integration

Our barcoding system uses two different landing pad types, hereafter referred to as type 1 and type 2. Both plasmids had a pUC origin and ampicillin resistance cassette in the vector backbone. The inserts into this 1998bp backbone were 6728bp and 6384bp, respectively, with ~450bp homology to the regions flanking the *CgTrp1* in the *HO* locus on either side. Between these flanking regions were modified versions of the *KanMX* and *CAN1* genes, as well as a *ccdB* gene that is toxic to sensitive *E. coli* strains. Many other components, including lox sites, artificial introns, and unexpressed *TRP1* genes, were also present in these plasmids, and the entirety of the annotated plasmids can be viewed in Supplementary Files 3 and 4. These extraneous elements – both in the plasmids and in our strain backgrounds – were included to enable capabilities that ultimately were not harnessed for the purposes of this study, such as mating, sporulation, and the inducible and selectable Cre-driven recombination of barcodes.

To generate diversely barcoded plasmid libraries, we cloned oligonucleotides containing random nucleotides into the type 1 and type 2 plasmids via a Golden Gate reaction (Engler, Kandzia, and Marillonnet 2008). This reaction replaced the *ccdB* gene in the plasmid. The barcoded plasmids were transformed via electroporation into *ccdB*-sensitive *E. coli*. Barcoded plasmids were then purified from these transformants using the Geneaid Presto^TM^ Mini Plasmid Kit (Cat. No. PDH300).

To barcode ancestral YCB137A and YCB140B strains, we took advantage of PmeI restriction endonuclease sites on either side of the *HO* homology regions of the plasmid, cutting and transforming (Gietz 2015) into the *HO* locus and replacing the *CgTRP1* gene.

To select for successful haploid yeast transformants, we used 200 μg/mL G418 (GoldBio, G-418), following up with a screen in SD-Trp (1.71 g/L Yeast Nitrogen Base Without Amino Acids and Ammonium Sulphate (Sigma-Aldrich, Y1251), 5 g/L ammonium sulfate (Sigma-Aldrich, A4418), 20 g/L dextrose (VWR #90000-904), 0.1 g/L L-glutamic acid (Sigma-Aldrich, G1251), 0.05 g/L L-phenylalanine (Sigma-Aldrich, P2126), 0.375 g/L L-serine (Sigma-Aldrich, S4500), 0.2 g/L L-threonine (Sigma-Aldrich, T8625), 0.01 g/L myo-Inositol (Sigma-Aldrich, I5125), 0.08 g/L adenine hemisulfate salt (Sigma-Aldrich, A9126), 0.035 g/L L-histidine (Sigma-Aldrich, H6034), g/L L-leucine (Sigma-Aldrich, L8000), 0.12 g/L L-lysine monohydrate (Acros Organics, CAS[39665-12-8]), 0.04 g/L L-methionine (Sigma-Aldrich, M9625), 0.04 g/L uracil (Sigma-Aldrich, U1128)). After ~25 generations of selection in liquid media, strains auxotrophic for tryptophan and resistant to G418 were arrayed into plates for experimental evolution.

Other successful transformants (of the same landing pad type) were mated to form diploids, which were selected for resistance to 300 μg/mL hygromycin B (GoldBio, H-270), 100 μg/mL nourseothricin sulfate (GoldBio, N-500), 200 μg/mL G418, and 1 mg/mL 5-fluoroorotic acid monohydrate (5-FOA) (Matrix Scientific, CAS[220141-70-8]) in S/MSG D media (1.71 g/L Yeast Nitrogen Base Without Amino Acids and Ammonium Sulphate, L-glutamic acid monosodium salt hydrate (Sigma-Aldrich, G1626), 20 g/L dextrose, 0.1 g/L L-glutamic acid, 0.05 g/L L-phenylalanine, 0.375 g/L L-serine, 0.2 g/L L-threonine, 0.01 g/L myo-Inositol, 0.08 g/L adenine hemisulfate salt, 0.035 g/L L-histidine, 0.11 g/L L-leucine, 0.12 g/L L-lysine monohydrate, 0.04 g/L L-methionine, 0.04 g/L uracil, 0.08 g/L L-tryptophan (Sigma-Aldrich, T0254)) for ~25 generations prior to arraying into 96-well plates alongside haploids for experimental evolution.

### Experimental evolution

Barcoded yeast were used to found 192 *MAT*a, 192 *MAT*α, and 162 diploid populations for evolution, respectively (though most haploid populations were excluded from further analysis due to the fixation of autodiploids). Each population was founded by a uniquely barcoded single colony or uniquely barcoded colonies that were then mated to form a diploid (see “Strain generation” section above), and was subsequently propagated in a well of an unshaken flat-bottom polypropylene 96-well plate in one of three conditions: YPD (1% Bacto yeast extract (VWR #90000–726), 2% Bacto peptone (VWR #90000–368), 2% dextrose) at 30°C, YPD at 37°C, and YPD+0.2% acetic acid (Sigma Aldrich #A6283) at 30°C (128 μL/well). Each 96-well plate contained diploid and haploid populations of both mating types (with each mating type occupying one side of the plate) and 5 empty wells to monitor for potential cross contamination. With the exception of the YPD at 37°C condition, the evolution conditions were arranged in a checkered pattern on each 96-well plate to minimize potential plate effects. Daily 1:2^10^ dilutions (bottleneck ~ 10^4^ cells) were performed using a Biomek-FX pipetting robot (Beckman-Coulter) after thorough resuspension by shaking on a Titramax 100 orbital plate shaker at 1,200 r.p.m. for at least 1 min. Populations underwent daily transfers for ~1000 generations (~10 generations/day); every 50 generations, populations were mixed with glycerol to a final concentration of 8% for long-term storage at −80°C. No contamination of blank wells was observed over the course of the evolution experiment. One of the 96-well plates was dropped at generation 170 and evolution was resumed by thawing and reviving populations from the generation 150 archive; thus, all future archives of populations on this plate lagged 40 generations behind the populations on all other plates.

### Nucleic acid staining for ploidy

Populations frozen at generation 1000 of the evolution experiment were thawed and revived by diluting 1:2^5^ in YPD. The following day, saturated cultures were diluted 1:10 into 120 μL of sterile water in round-bottom polystyrene 96-well plates. Plates were centrifuged at 3,000xg for 3 minutes, the supernatant was removed, and cultures were resuspended in 50 μL sterile water. 100 μL of ethanol was added to each well, the cultures were mixed thoroughly and placed at 4°C overnight. The following day, the cultures were centrifuged, the ethanol solution was removed, and 65 μL RNase A (VWR #97062-172) solution (2 mg/mL RNase A in 10 mM Tris-HCl, pH 8.0 + 15 mM NaCl) was added to each well and the cultures were incubated at 37°C for 2 h. Then 65 μL of 300 nM SYTOX green (Thermo Fisher Scientific, S-34860) was added to each well and the cultures were mixed and incubated at room temperature in the dark for 30 min. Fluorescence was measured by flow cytometry on a BD LSRFortessa using the FITC channel (488 nm). Ploidy was assessed by comparing the fluorescence distributions of evolved populations to known haploid and diploid controls of the same strain. By generation 1000, all 192 *MAT*a populations had autodiploidized, and 172 of the *MAT*α populations had autodiploidized, as judged by the absence of a clear haploid peak. Only the remaining 20 haploid *MAT*α populations were included in the bulk fitness assays described below.

### Bulk fitness assays

Populations frozen at generations 0, 200, 400, 600, 800, and 1000 of the evolution experiment were thawed by diluting 1:2^5^ in YPD. The following day, once these cultures had grown to saturation, equivalent volumes of each population were pooled by ploidy for each generation (12 pools total). For the haploid populations, evolved populations were only pooled if they were verified to be haploid at the end of the evolution experiment (see “Nucleic acid staining for ploidy” section above). Each of the haploid pools was spiked with 5 uniquely barcoded ancestral reference strains of the same mating type at 4X the volume of each evolved population; each of the diploid pools was spiked with 10 reference strains at 4X the volume of each evolved population. The resulting pools comprised time point zero for the bulk fitness assay (BFA) and were diluted 1:2^10^ in the appropriate media (described below) and divided between 16 wells (128 μL/well) of flat-bottom polypropylene 96-well plates. The BFA was performed in each of the three evolution environments (YPD at 30°C, YPD at 37°C, and YPD+0.2% acetic acid at 30°C), in addition to two novel environments (YPD at 21°C and YPD+0.4M NaCl at 30°C). The 16 wells of each pool comprised two technical replicates of 8 wells. Every 24 hours (or every 48 hours in the case of the YPD 21°C environment) the populations were resuspended by shaking on a Titramax 100 orbital plate shaker at 1,200 r.p.m. for at least 1 min and the contents of the 8 wells constituting each replicate were combined, mixed, and diluted 1:2^10^ into 8 new wells using a Biomek-FX pipetting robot (Beckman-Coulter). This split-pool strategy was designed to mimic the evolution conditions while maintaining sufficient diversity for bulk fitness measurements. At BFA timepoints 0, 10, 30, and 50 generations, 1 mL of the diploid pool was combined with 200 μL of the haploid pool for each generation, this culture was centrifuged at 21,000 × g for 1 minute, the supernatant was removed, and the pellet was stored at −20°C for downstream DNA extraction and sequencing.

### Sequencing library preparation

Genomic DNA was extracted from cell pellets using zymolyase-mediated cell lysis (5 mg/mL Zymolyase 20T (Nacalai Tesque), 1 M sorbitol, 100 mM sodium phosphate pH 7.4, 10 mM EDTA, 0.5% 3-(N,N-Dimethylmyristylammonio)propanesulfonate (Sigma T7763), 200 μg/mL RNase A, 20 mM DTT), binding on silica columns (IBI scientific, IB47207) with 4 volumes of guanidine thiocyanate (4.125 M guanidine thiocyanate, 100 mM MES pH 5, 25% isopropanol, 10 mM EDTA), washing with wash buffer 1 (10% guanidine thiocyanate, 25% isopropyl alcohol, 10 mM EDTA) and wash buffer 2 (20mM Tris-HCl pH 7.5, 80% ethanol), and eluting in 50 μL 10 mM Tris pH 8.5, as previously described (Nguyen Ba et al. 2019). Two rounds of PCR were performed to generate amplicon sequencing libraries for sequencing the barcode locus. In the first round of PCR, the barcode locus was amplified with primers containing unique molecular identifiers (UMI), generation-specific inline indices, and partial Illumina adapters (see Supplementary File 5 for primer sequences). This 20 μL 10-cycle PCR reaction was performed using Q5 polymerase (NEB M0491L) following the manufacturer’s guidelines, using 10 μL (~250 ng) of gDNA as template, annealing at 54°C, and extending for 45 seconds. The first-round PCR products were then purified using one equivalent volume of DNA-binding beads (Aline Biosciences PCRCleanDX C-1003-5) and eluting in 33 μL 10 mM Tris pH 8.5. In the second-round PCR, the remainder of the Illumina adapters and sample-specific Illumina indices were appended to the first-round PCR products (see Supplementary Table 5 for primer sequences). The second round PCR was performed using Kapa HiFi HotStart polymerase (Kapa Bio KK2502) following the manufacturer’s guidelines for a 25 μL reaction, using 17.25 μL of first round PCR product, annealing at 63°C and extending for 30 seconds for 26 cycles. The second-round PCR products were then purified using one equivalent volume of DNA-binding beads and eluting in 33 μL 10 mM Tris pH 8.5. Following bead cleanup, the concentration of the PCR products was quantified using the Accugreen High Sensitivity dsDNA Quantitation Kit (Biotium 31068). Sequencing libraries were then pooled equally and sequenced on a NextSeq500 Mid flow cell (150 bp single-end reads).

### BFA barcode enumeration and fitness inference

Lineage fitnesses were inferred from the concatenated sequencing data yielded by two separate NextSeq500 Mid flow cells (150bp single-end reads). The second of these two runs allowed for deeper sequencing of specific BFA timepoints to enable superior determination of barcode frequencies associated with less fit lineages in certain environments. The second run also allowed sequencing of libraries that were omitted from the first run.

Once fastq files were concatenated, barcode information was extracted as described below. However, in addition to subjecting the barcode regions to error-tolerant ‘fuzzy’ matching based on regular expressions, we allowed for fuzzy matching of the epoch-specific inline indices. For the indices, we applied a list of decreasingly strict regular expressions, looking for exact matches, then 1 mismatch, then 2 mismatches. For the indices associated with epochs 6, 8, and 10, which were longer than the indices associated with epochs 0, 2, and 4, we allowed up to 3 mismatches.

Then, as with the barcode association mapping, we used a previously described “deletion-error-correction” algorithm (Johnson et al. 2019) to correct errors in barcode sequences induced by library preparation and sequencing.

To check for cross-contamination between wells during library preparation and index-hopping during sequencing, we searched for reads where the inline index was inconsistent with the associated pairs of Illumina indices. In almost all cases, we found little evidence of cross contamination (<< 1%). In one case, corresponding to landing pad type 2 of the 30°C replicate 2 BFA 10-generation timepoint for generation-1000 populations, we found that 11,484 of the 258,462 reads (4.4%) included the inline index associated with the generation-200 populations. We removed all apparently cross-contaminating reads from our analysis.

Then, we summed reads associated with all barcodes in a given population, since some populations contained more than one unique barcode (or, in the case of diploids, more than two unique barcodes). In addition, some barcodes were present in the BFAs that could not be confidently assigned to a single well, representing 0.3% of all reads. These were summed together and retained in the dataset.

To determine the fitness of each population over time and across environments and technical replicates, we measured the log-frequency slope for each population in two intervals: between assay timepoints 10 and 30 and between timepoints 30 and 50 generations. Frequencies were calculated separately for each landing pad type. We scaled these values of fitness (*s*) by subtracting out the corresponding median log-frequency slope of a set of between 2 and 5 reference ancestral populations of each ploidy and landing pad type, which were included in every BFA to allow comparisons of fitness across the evolutionary time course. The source data file indicates these reference populations. For a given BFA and interval, *s* values only were calculated this way if the mean number of reads for the reference populations was greater than 5. If not, these intervals were excluded from subsequent analysis.

To determine *s* values for each population in each environment at each generation, interval-specific *s* estimates were averaged. Then, *s* estimates from each of the two technical replicates were averaged, producing a final *s* estimate. The standard error of this final *s* estimate was calculated from the two technical replicate *s* estimates.

To clarify our downstream analyses, we excluded 19 outlier diploid populations whose ancestors differed from the mean ancestral fitness by at least 4% in at least one environment. We believe we see such divergent ancestral fitness values due mutations that emerged during the process of selecting colonies, mating, and performing purifying selection for ~50 generations on barcoded transformants immediately prior to evolution.

To account for the offset in plate 2 progress through evolution, plate 2 population fitness estimates for 200, 400, 600, and 800 generations were linearly interpolated from fitnesses on either side, e.g., gen 200 fitness inferred from gen 160 and gen 360 fitnesses. Fitness estimates for gen 1000 were extrapolated linearly from gen 760 and gen 960 fitnesses. The standard error of the *s* estimate for gen 160 was used for gen 200 fitness, the standard error of *s* for gen 360 was used for gen 400 fitness, and so on.

### Barcode association

To map barcodes to wells of the evolution experiment, we pooled ancestral strains in equal volumes from across the eight evolution plates, creating three sets of pools: column-specific pools (n=12), row-specific pools (n=8), and plate-specific pools (n=8). We then lysed portions of these pools by diluting in yeast lysis buffer (1mg/mL Zymolyase 20T, 0.1M Sodium phosphate buffer pH 7.4, 1M sorbitol, 10 mg/mL SB3-14 (3-(N,N-Dimethylmyristylammonio)propanesulfonate (Sigma T7763)) at 37°C for 1hr and 95°C for 10min. Two rounds of PCR were then performed to generate amplicon sequencing libraries for sequencing the barcode locus (both landing pad versions). In the first round, the barcode locus was amplified via a 10-cycle PCR reaction with Kapa HiFi HotStart polymerase (Kapa Bio KK2502), annealing at 58°C for 30 s and extending at 72°C for 30 s, with a final 10 min extension. PCR products were then purified using one equivalent volume of DNA-binding beads and eluting in 20 μL water. Following bead purification, a second-round PCR reaction was performed using 1.5 μL of each of a unique pair of Illumina indices (see Supplementary File 5 for primer sequences) with Kapa HiFi Hotstart ReadyMix (2X) in a 15 μL reaction, with 4.5 μL of first-round PCR product as template, annealing at 61°C and extending for 30 seconds for 30 cycles. The second-round PCR products were then purified using 0.8x DNA-binding beads (Aline Biosciences PCRClean DX C-1003-5), washed 2x with 80% ethanol and eluted in 50 μL of molecular biology-grade water. Following bead cleanup, the concentration of the second round PCR products was quantified using the Accugreen High Sensitivity dsDNA Quantitation Kit (Biotium 31068). These libraries were then normalized, pooled, and sequenced on a NextSeq500 High flow cell (150 bp paired-end reads).

To extract barcode information from sequencing reads, we followed Johnson et al. (2019), using a list of decreasingly strict regular expressions (using the python regex module https://pypi.org/project/regex/). For landing pad 1, this was:

’(TCTGCC)(\D{22})(CGCTGA)’,
’(TCTGCC)(\D{20,24})(CGCTGA)’,
’(TCTGCC){e<=1}(\D{22})(CGCTGA){e<=1}’,
’(TCTGCC){e<=1}(\D{20,24})(CGCTGA){e<=1}’

For landing pad 2, this was:

’(TCTCTG)(\D{22})(AGTAGA)’,
’(TCTCTG)(\D{20,24})(AGTAGA)’,
’(TCTCTG){e<=1}(\D{22})(AGTAGA){e<=1}’,
’(TCTCTG){e<=1}(\D{20,24})(AGTAGA){e<=1}’

Then, after parsing and tallying barcodes in each sequencing library, we used the “deletion-error-correction” algorithm described by Johnson et al. (2019) to correct errors in barcode sequences induced by library preparation and sequencing.

To triangulate the position of each barcode across the eight plates, for each error-corrected barcode that appeared in the sequencing data, we tabulated which barcodes were present in which libraries, and how many reads were associated with each barcode in each library. These data allowed us to determine the wells in which barcodes belonged.

### IQR variability analysis

Fitness variability was examined by plotting box-and-whisker plots of population mean fitness values, where the line, box, and whiskers represent the median, quartiles, and data within 1.5xIQR of each quartile, respectively, and outlier populations beyond whiskers are shown as points (**Fig. 5A**). To compare the resulting IQR for various evolution conditions and fitness assay environments, 95% confidence intervals of the IQR were calculated from bootstrapped interval-specific replicate s measurements (**Fig. 5B**).

To evaluate whether home environment fitness variance was less than away environment fitness variances at each evolution timepoint, we applied a Brown-Forsythe test (Brown and Forsythe 1974). Since this test is typically a two-tailed test, and we wanted instead to employ a one-tailed test, we used the *z* scores from the Brown-Forsythe test to arrive at a two-tailed *t*-statistic. We could then obtain a one-tailed *p*-value with this *t*-statistic, evaluated at *N* − 1 degrees of freedom, where *N* is the number of populations in consideration.

### Principal components analysis

All principal components analysis excluded ancestral reference populations. To minimize the influence of varying scales of data features on the analysis, fitness values for each field – corresponding to fitness in a given assay environment, possibly at a specific evolutionary timepoint – were standardized to have a mean of 0 and standard deviation of 1 using the scikit-learn StandardScaler function. We then used the scikit-learn PCA() function.

### Clustering metric

To quantify the degree of clustering by evolution condition in the 2-dimensional principal component analyses, the NearestNeighbors algorithm in the scikit-learn python package was implemented to identify the five nearest neighbors for each population in the 2-dimensional PC1 versus PC2 plots (Figs. 4A,B). The clustering metric plotted in **Fig. 4D** is the number of five nearest neighbors that belong to the same evolution condition as the focal population, averaged for each evolution condition. Error bars represent 95% confidence intervals of the mean clustering metric, which were calculated by performing the PCA and clustering analysis on bootstrapped interval-specific replicate *s* measurements. The null expectation for populations to cluster by evolution condition was computed by permuting the evolution condition 1000 times and performing the clustering analysis as described above. The permuted clustering metrics were then compared to the true mean clustering metric by a two-sided Student’s *t*-test (using the Scipy.stats ttest_ind_from_stats function).

### Non-monotonicity analysis

To assess non-monotonicity, we linearly interpolated fitness at 500 generations for each population in each assay environment. We achieved the interpolated standard errors in fitness by taking the square root of the sum of the squares of the errors associated with the fitnesses used in the interpolation and dividing by two. For evolution plate 2 populations, which were offset from the others by 40 generations, we took a weighted average for the interpolation (500 generation fitness estimate) and extrapolation (1000 generation fitness estimate) steps. For the 500 generation fitness standard error estimate, we adapted this weighting approach for the standard error propagation as described for the other populations. For the 1000 generation fitness standard error estimate, we used the error assigned to the generation 960 fitness estimate. Then, we calculated the change in fitness (Δ*s*) between 0 and 500 generations and between 500 and 1000 generations for each population in each environment. The standard errors of these Δs were the square root of the sum of the squares of the two fitnesses used in the calculation. Finally, we plotted these Δ*s* values as x-y coordinates. If a point and its error bars were completely within the top-left or lower-right quadrant -- corresponding to an increase followed by a decrease, or a decrease followed by an increase, over the 1000-generation experiment -- these were considered to be “clearly non-monotonic.” We applied a χ2 test to evaluate the significance in the difference in the frequency of non-monotonicity in home versus away trajectories.

## ACKNOWLEDGMENTS

We thank Parris T. Humphrey for assistance with experimental design and experimental protocols, and we thank Anurag Limdi for help with strain construction. We also thank Milo S. Johnson for helpful comments on the manuscript. C.B. acknowledges the support of the Department of Defense (DoD) through the National Defense Science & Engineering Graduate (NDSEG) Fellowship Program, as well as NIH training grant support (Joint Training Program in Molecules, Cells and Organisms, T32 Grant #GM007598). A.M.P. acknowledges support from the Howard Hughes Medical Institute Hanna H. Gray Postdoctoral Fellowship Program. M.M.D. acknowledges support from grant PHY-1914916 from the NSF and grant GM104239 from the NIH. The computations in this paper were run on the FASRC Cannon cluster supported by the FAS Division of Science Research Computing Group at Harvard University.

## ADDITIONAL FILES

### Supplementary files

- Supplementary file 1. Animation of Figure 3.
- Supplementary file 2. Strain creation tables.
- Supplementary file 3. Plasmid for landing pad 1 barcode integration.
- Supplementary file 4. Plasmid for landing pad 2 barcode integration.
- Supplementary file 5. Primers used in this study.
- Figure 2 – source data 1. Bulk fitness assay read counts and measured fitnesses.
- Figure 4 – source data 1. Principal component analyses presented in Figure 4A.
- Figure 4 – source data 2. Principal component analysis presented in Figure 4B.
- Figure 4 – source data 3. Principal component analyses presented in Figure 4 – figure supplement 1A.
- Figure 4 – source data 4. Principal component analysis presented in Figure 4 – figure supplement 1B.
- Transparent reporting form

## DATA AVAILABILITY

All the strains used here are available from the corresponding author upon request. Raw amplicon sequencing reads have been deposited in the NCBI BioProject database with accession number PRJNA739738. Source data files are listed in appropriate figure legends. Analysis code is available at https://github.com/amphilli/pleiotropy-dynamics.

---

**Figure 1–figure supplement 1.**
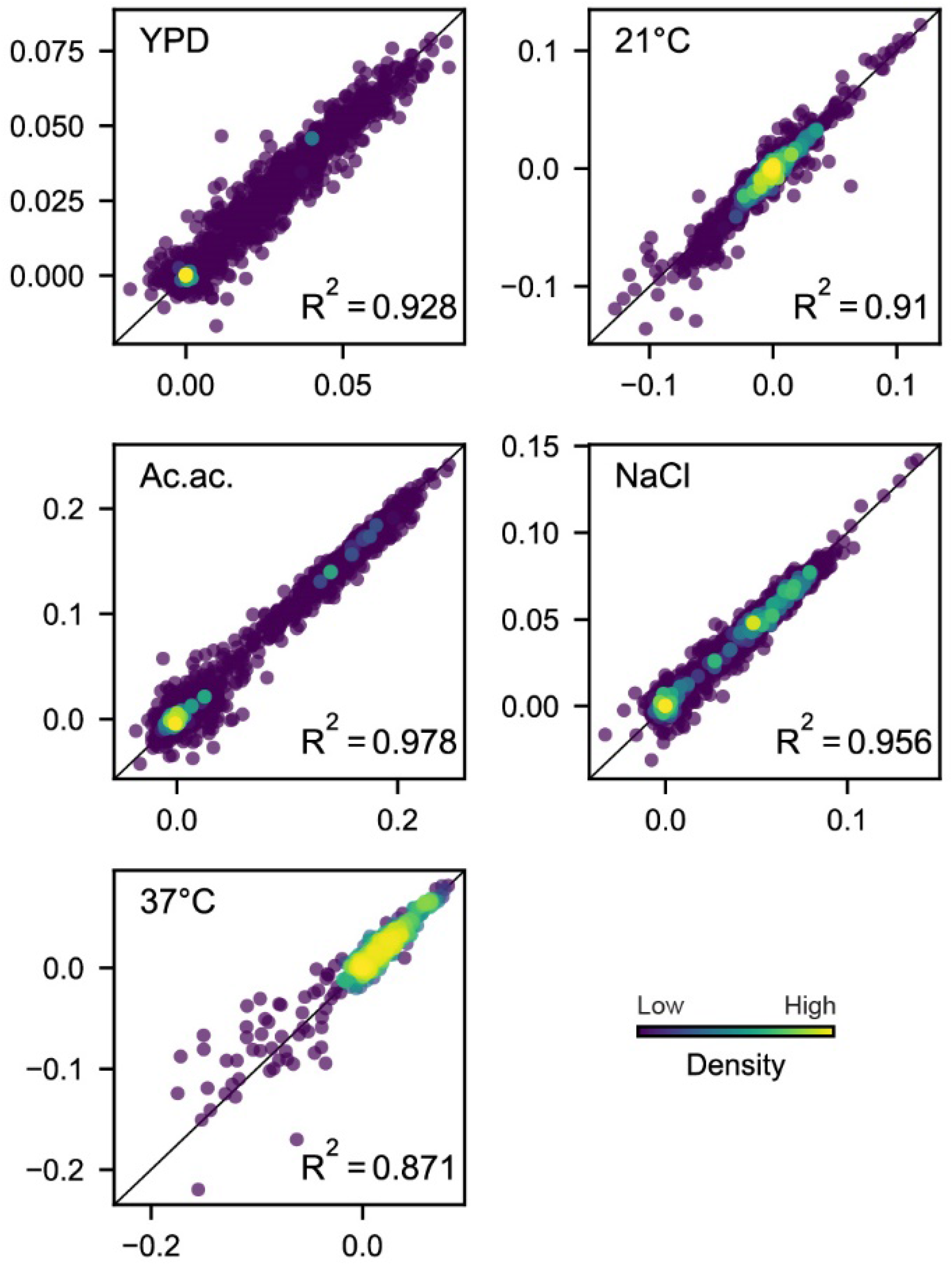
Comparison of technical replicate fitness measurements. Each dot corresponds to the fitness of a population at a given evolution timepoint in the environment indicated. Point color corresponds to the relative density of points, as determined by distance to five nearest points. The black line in each plot indicates x=y.

**Figure 2–figure supplement 1.**
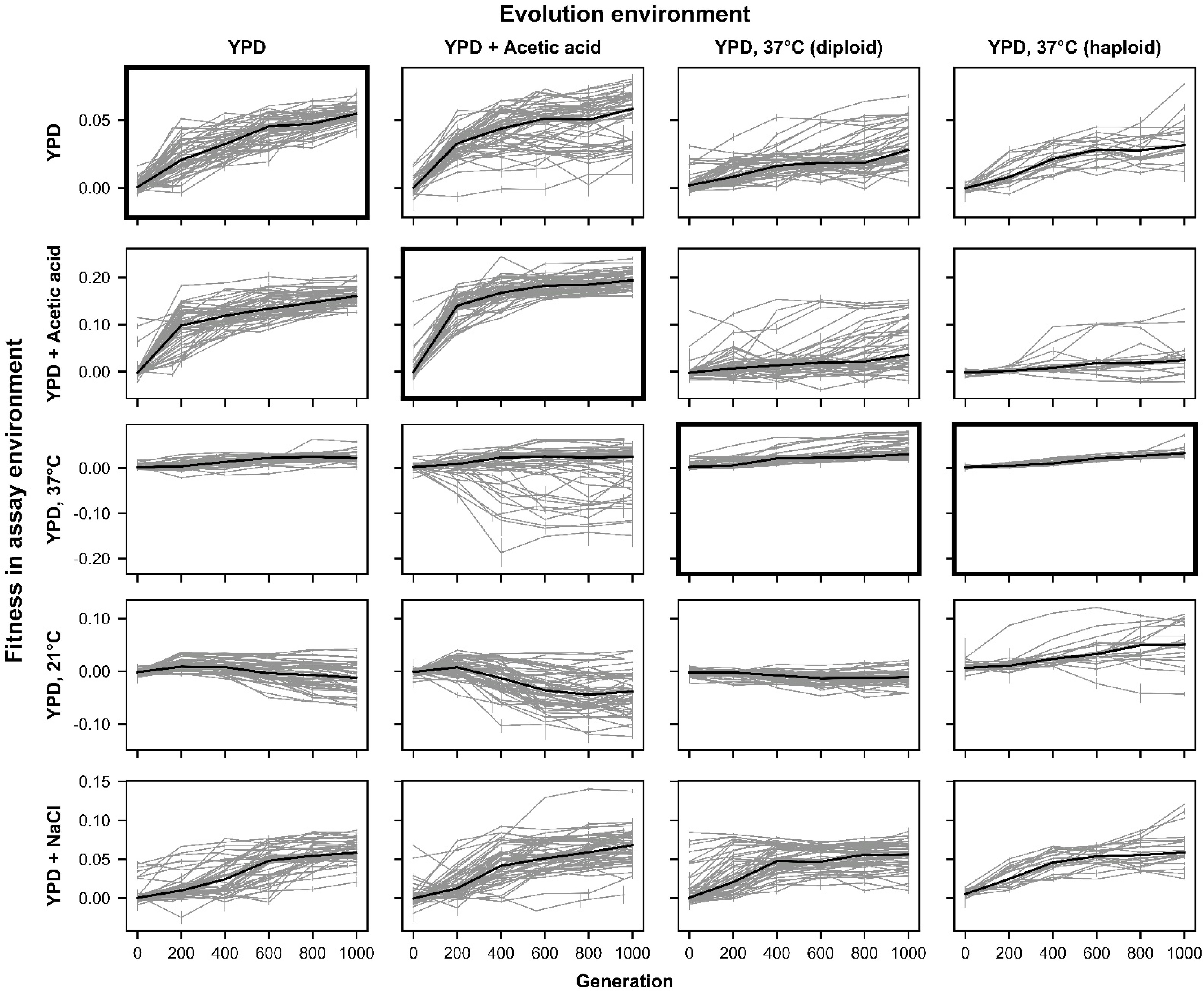
Fitness changes over 1000 generations of evolution for unfiltered data. Replicate populations for each evolution condition are shown in each column. Environments in which these populations’ fitnesses were assayed are shown in the rows. Plots for which evolution and assay environment are the same are indicated by a bold outer border. The black line in each plot indicates the median fitness. Error bars indicate standard error of the mean.

**Figure 3–figure supplement 1.**
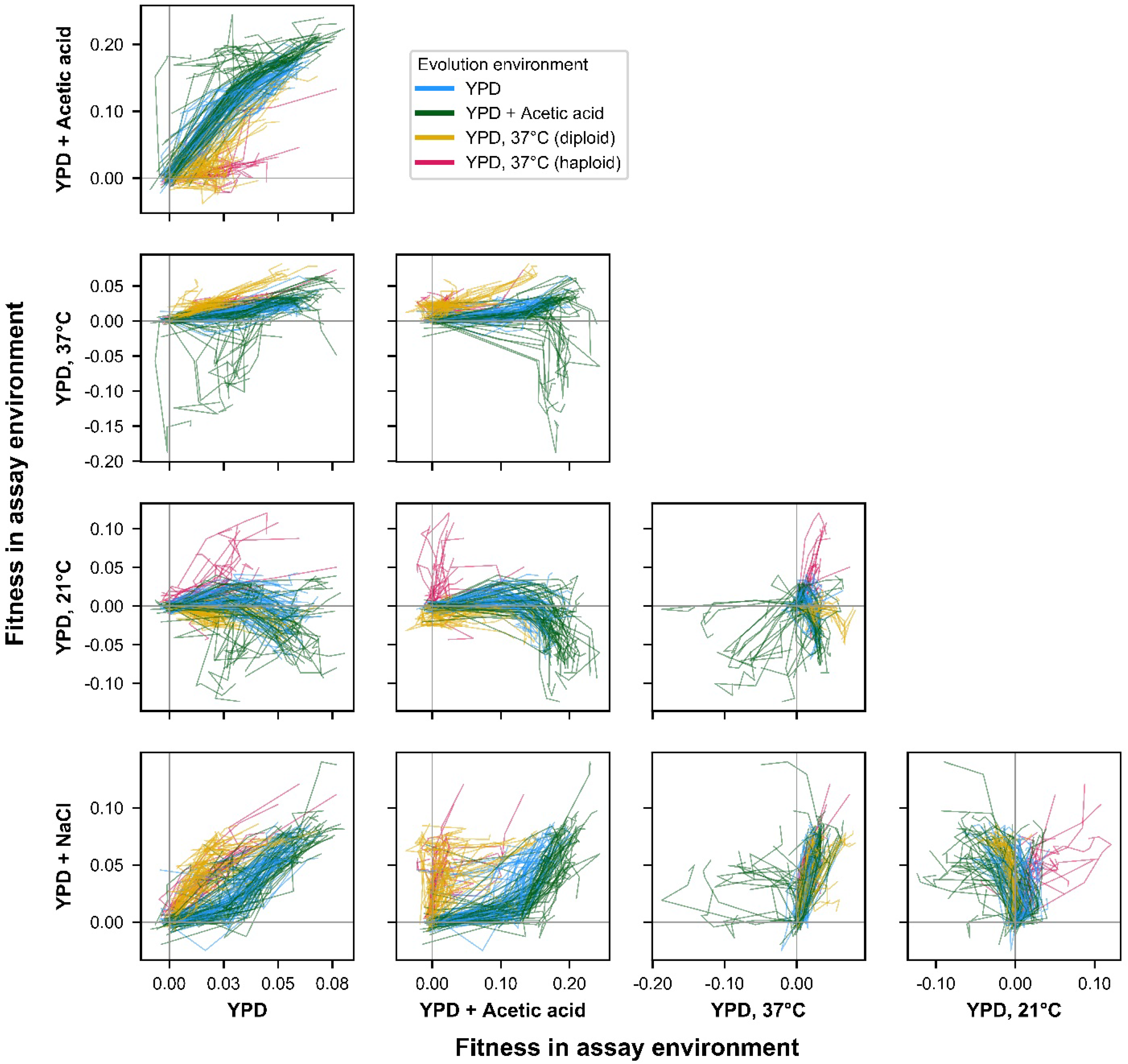
ExE evolutionary trajectories over 1000 generations of evolution in a constant environment for unfiltered data. Axes correspond to fitness in the indicated assay environments. Colors correspond to evolution condition. Grey vertical and horizontal lines indicate zero fitness relative to an ancestral reference in each environment.

**Figure 4–figure supplement 1.**
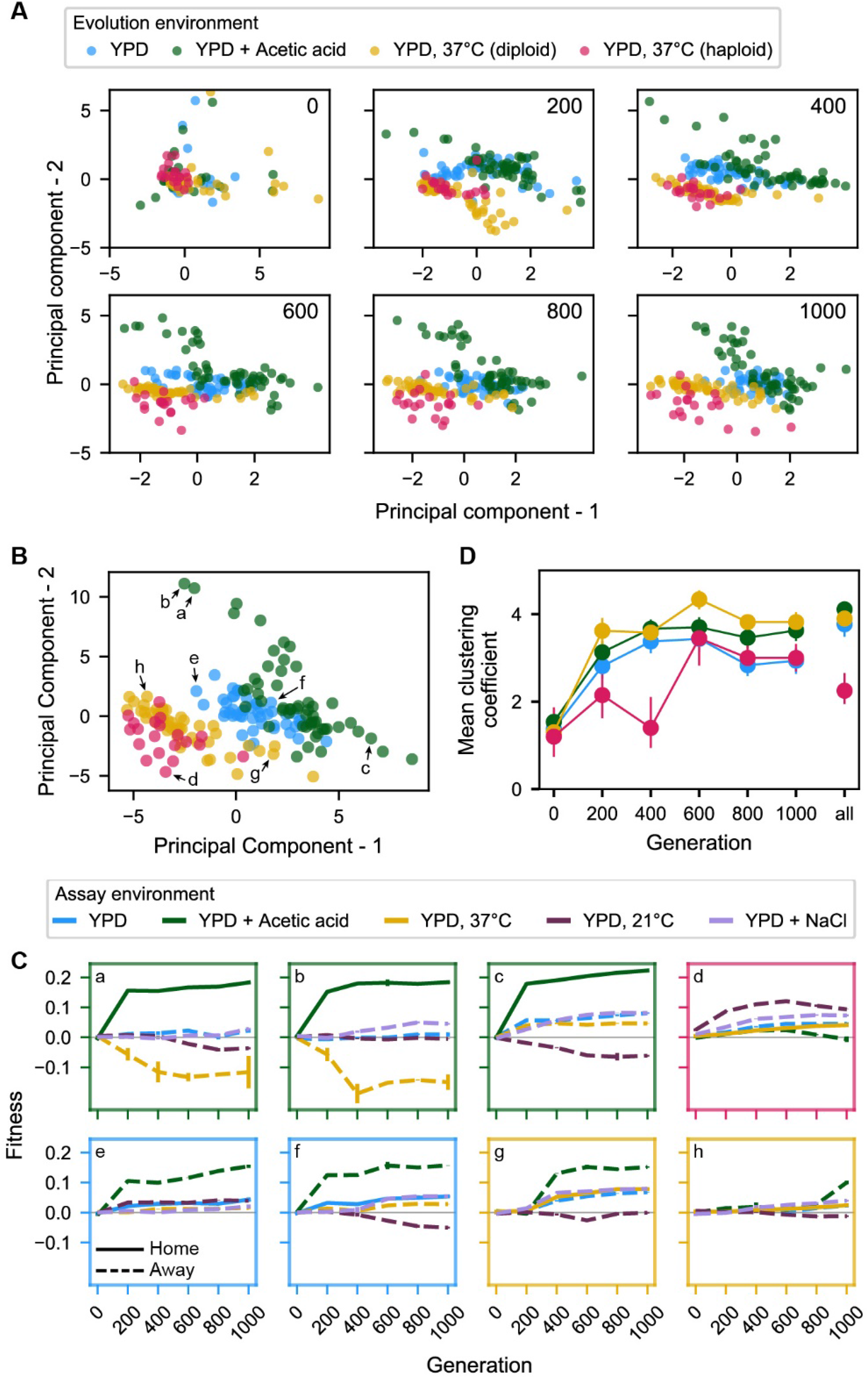
Principal component analysis of pleiotropy. (A-D) correspond to the same panels of Figure 4, except with analyses performed on the whole dataset including outlier populations. (C) is identical to Figure 4C.

**Figure 4 – figure supplement 2.**
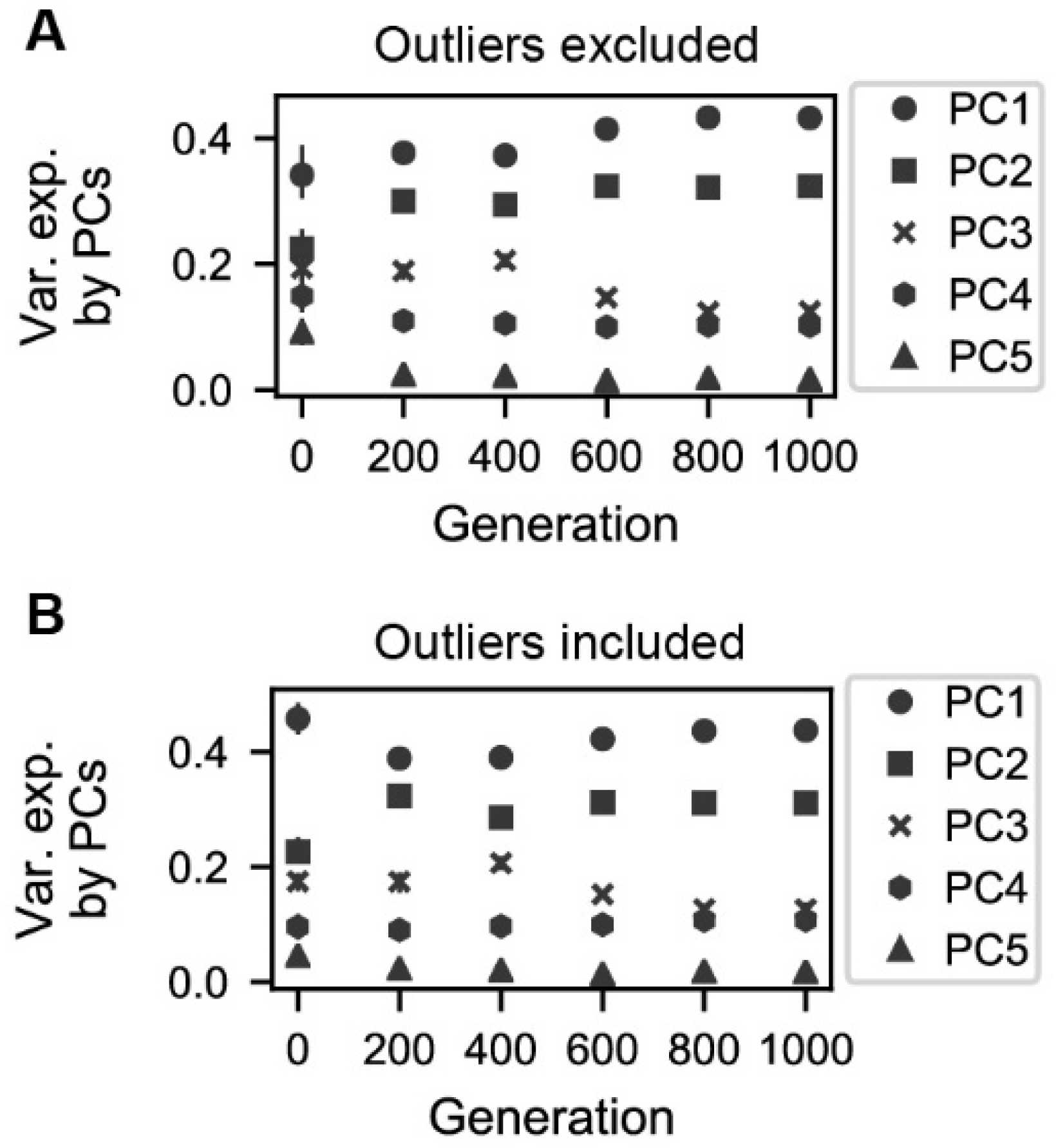
Variation explained by principal components. **(A)** Variance explained by five principal components corresponding to the PCAs conducted for each generation interval in Figure 4A. **(B)** Variance explained by five principal components corresponding to the PCAs conducted for each generation interval in Figure 4 – figure supplement 1A.

**Figure 4 – figure supplement 3.**
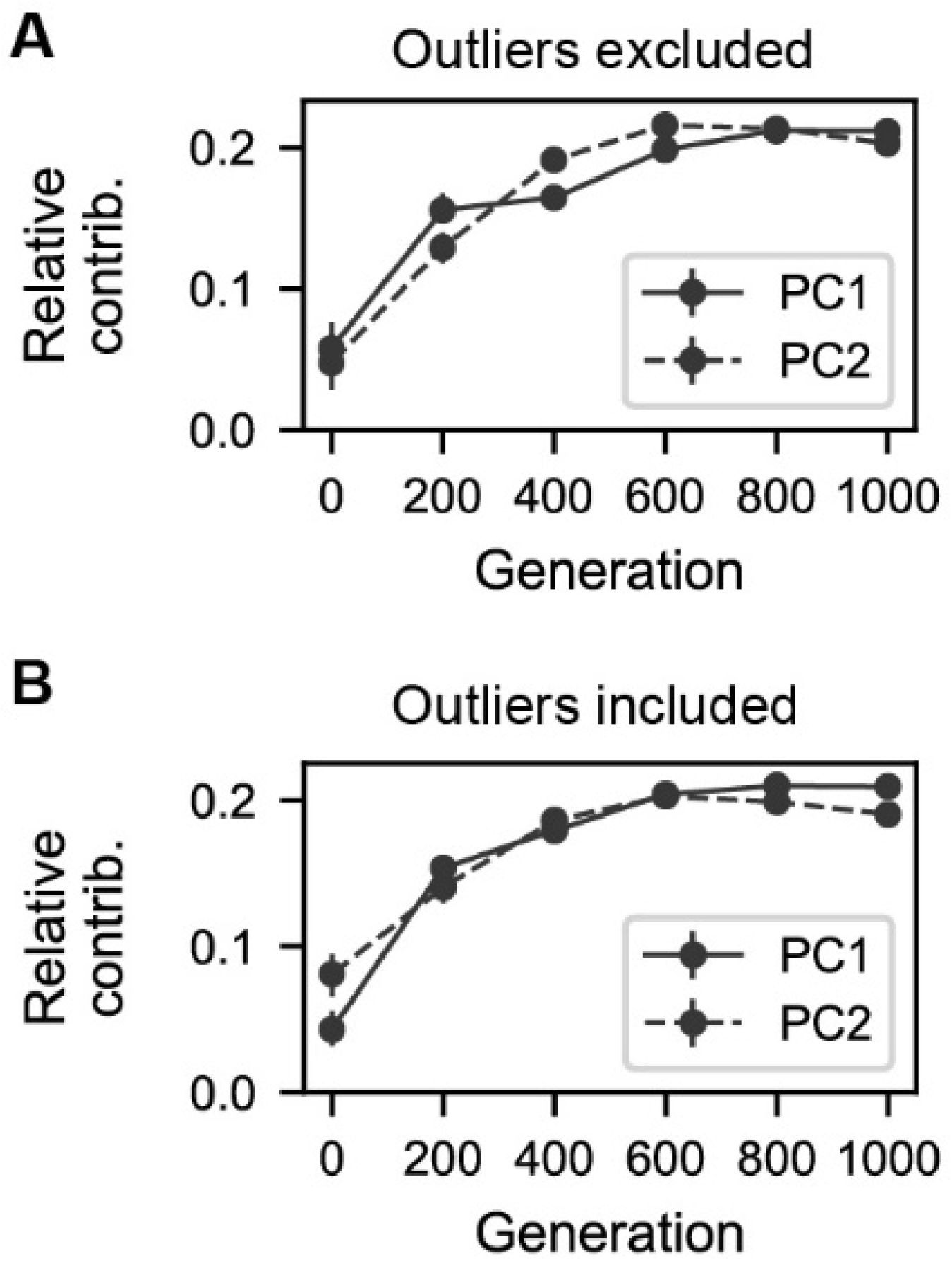
Contributions of generation intervals to principal components. **(A)** Summed magnitudes of contributions of assay environments at each interval to the two principal components presented in Figure 4B. **(B)** Summed magnitudes of contributions of assay environments at each interval to the two principal components presented in Figure 4 – figure supplement 1B.

**Figure 4 – figure supplement 4.**
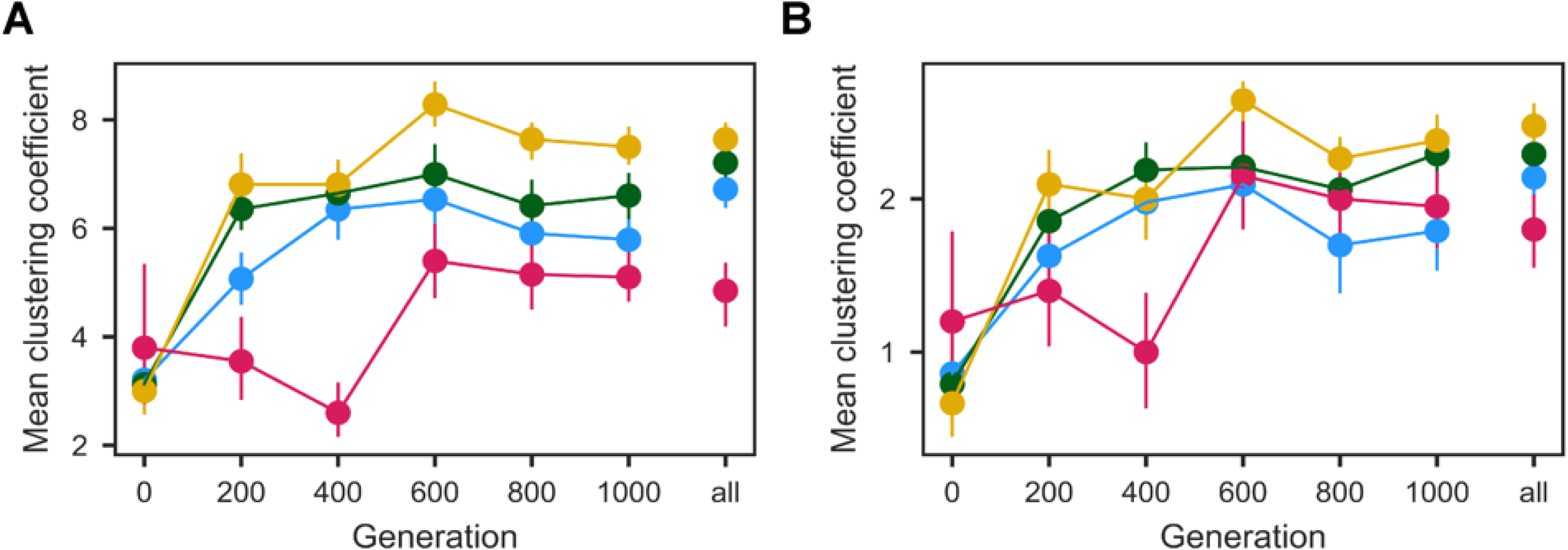
Population clustering in PCA as in Figure 4D quantified for **(A)** ten and **(B)** three nearest neighbors. Clustering metrics were averaged for each evolution condition to calculate point estimates; error bars represent 95% confidence intervals of the mean clustering metric, estimated by performing PCA on bootstrapped replicate fitness measurements.

**Figure 5–figure supplement 1.**
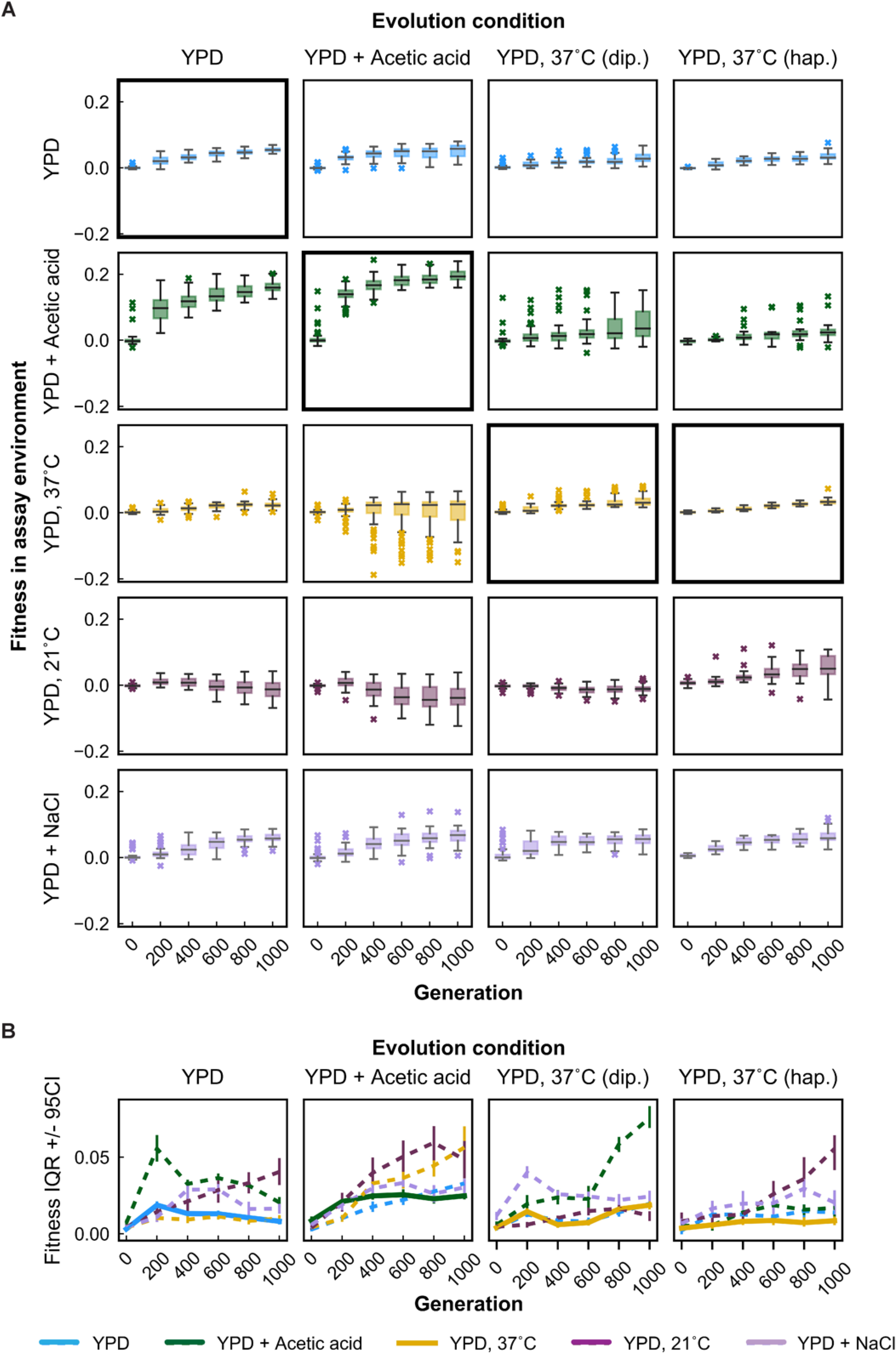
Variability in fitness over time for unfiltered data. **(A)** Box plots summarizing population mean fitness over time for each evolution condition (columns) in each assay environment (rows). Line, box, and whiskers represent the median, quartiles, and data within 1.5xIQR of each quartile, respectively; outlier populations beyond whiskers are shown as points. **(B)** IQR from box plots in (A) are plotted as a function of time for each evolution condition and assay environment. IQR for fitness measured in home and away environments are represented by solid and dashed lines, respectively. Error bars represent 95% confidence intervals of IQR calculated from bootstrapped replicate fitness measurements.

**Figure 5–figure supplement 2.**
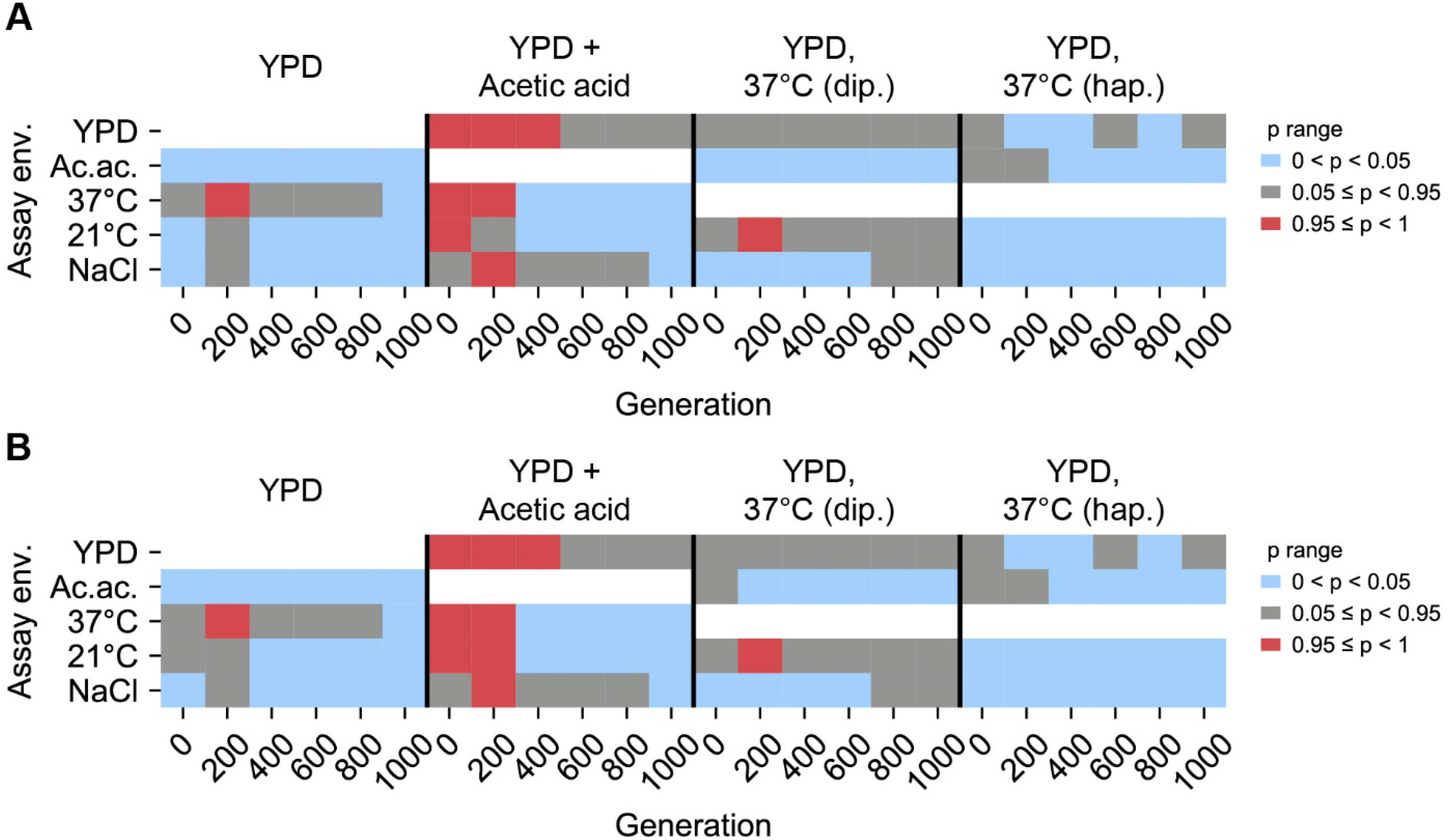
Statistical test of difference in variance between home, away environments. Brown-Forsythe test *p* values for paired comparisons of fitness variance in home environment and away environment for populations evolved in each evolution condition (columns). White boxes correspond to invalid self-comparisons. *p* values represent a one-sided test in which the alternative hypothesis is that home variance is less than away variance. 0 < *p* < 0.05 (blue) indicates home variance significantly less than away variance. 0.95 ≤ *p* < 1 (red) indicates home variance significantly greater than away variance. **(A)** Excluding outliers. **(B)** Including outliers.

**Figure 6–figure supplement 1.**
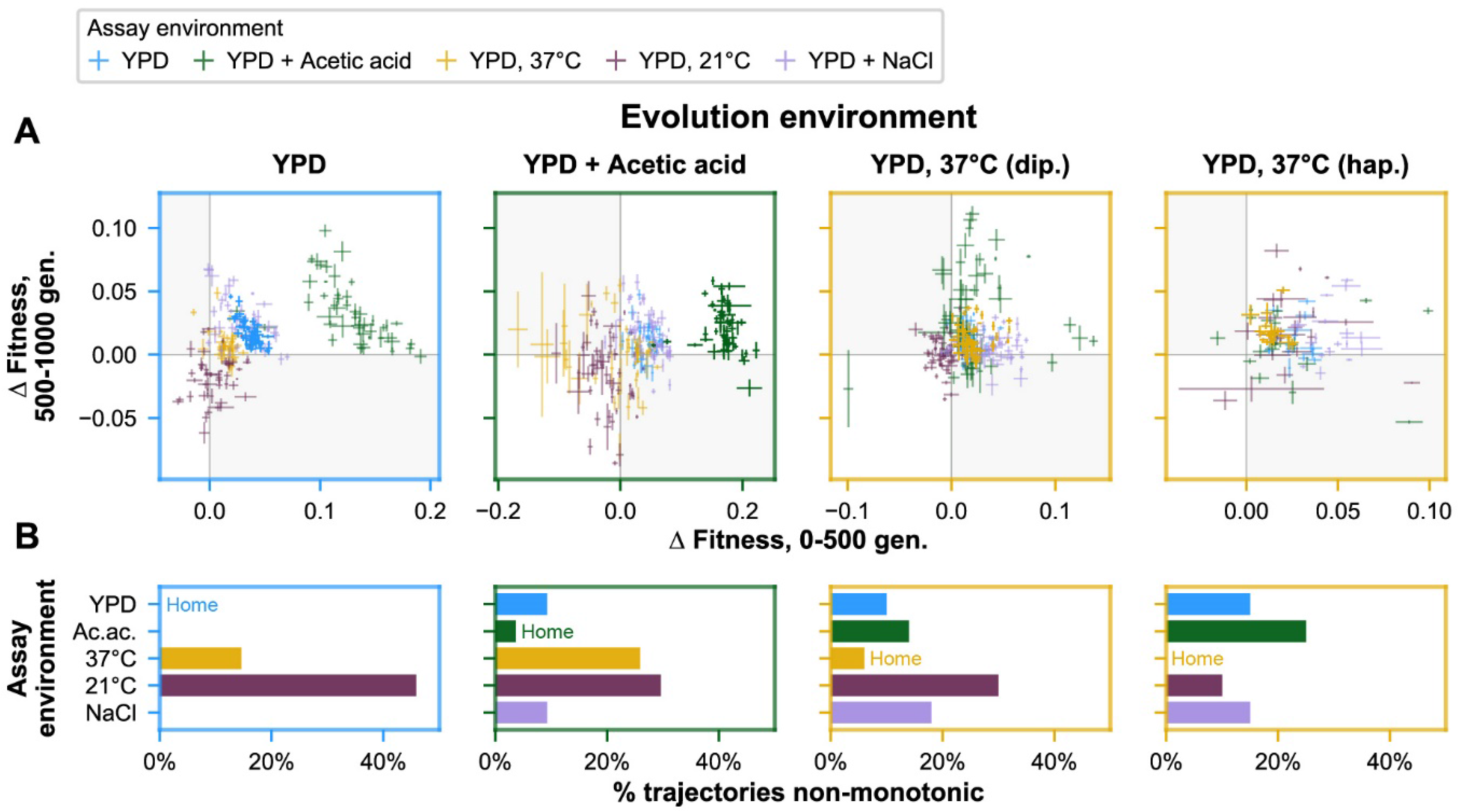
Non-monotonicity in evolutionary trajectories for unfiltered data (outliers included). **(A)** Each panel shows—for each of the 5 assay environments—the change in fitness over the first 500 (x-axis) and second 500 (y-axis) generations of evolution of each population in a given evolution environment. Populations that fall in shaded quadrants have trajectories that are non-monotonic. Points corresponding to fitness in the home environment are colored more opaquely than points corresponding to fitness in away environments, and panel borders have been colored to match the home environment. Fitness at generation 500 has been interpolated. **(B)** Each panel corresponds to a given evolution environment and shows the proportion of populations evolved in that environment that exhibit clearly non-monotonic fitness trajectories in (A). “Clearly non-monotonic” trajectories are those populations (points) in (A) that fall in the grey quadrants and whose error bars (1 standard error in either direction) do *not* span either the x- or y-axis. As in (A), bars corresponding to the home environment are colored more opaquely than bars corresponding to away environments. As with the outliers-excluded data, populations exhibit clearly non-monotonic trajectories in away environments much more commonly than in home environments (*p* < 0.0001), with most of these reflecting initially positive pleiotropic effects.

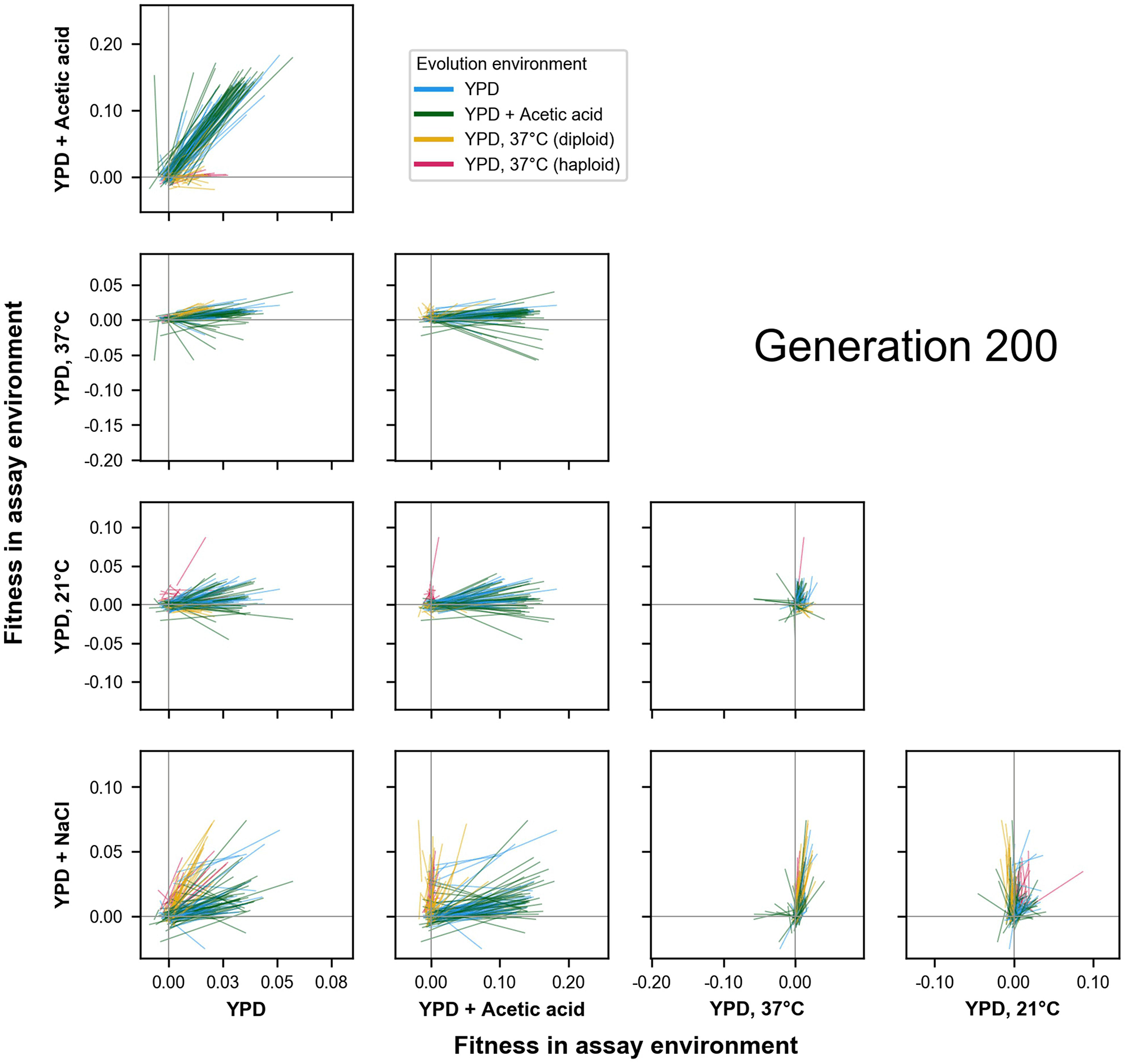

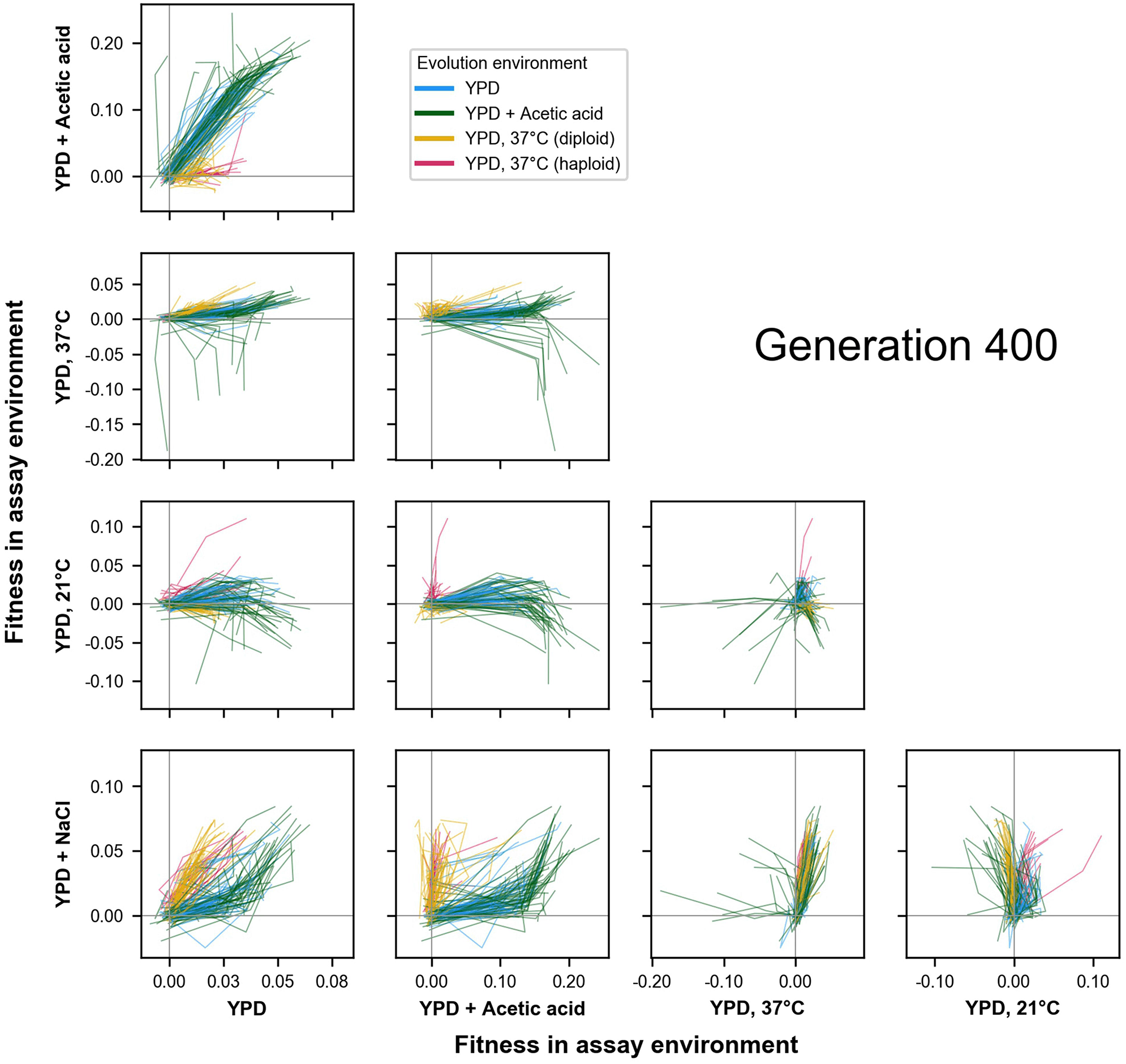

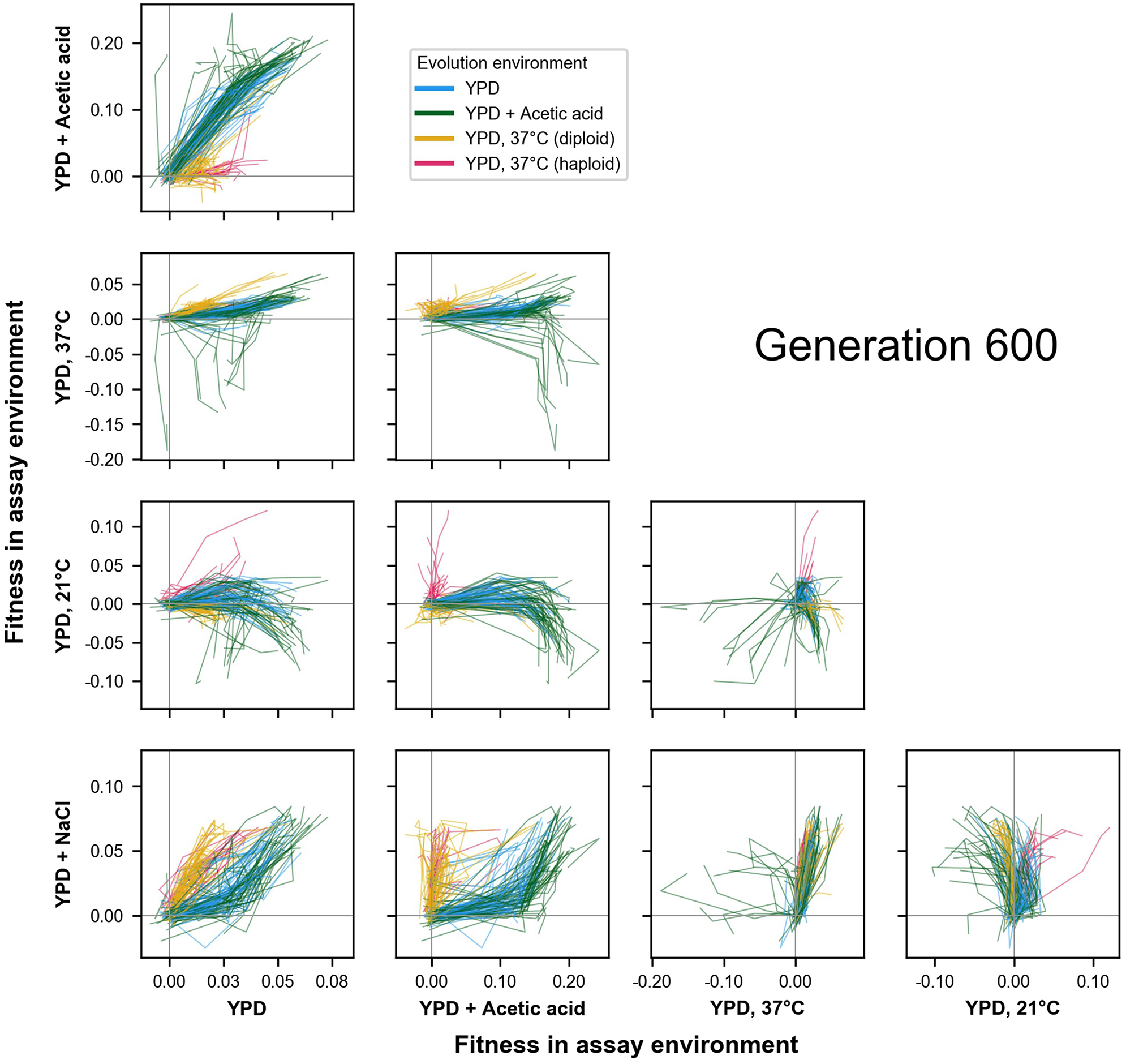

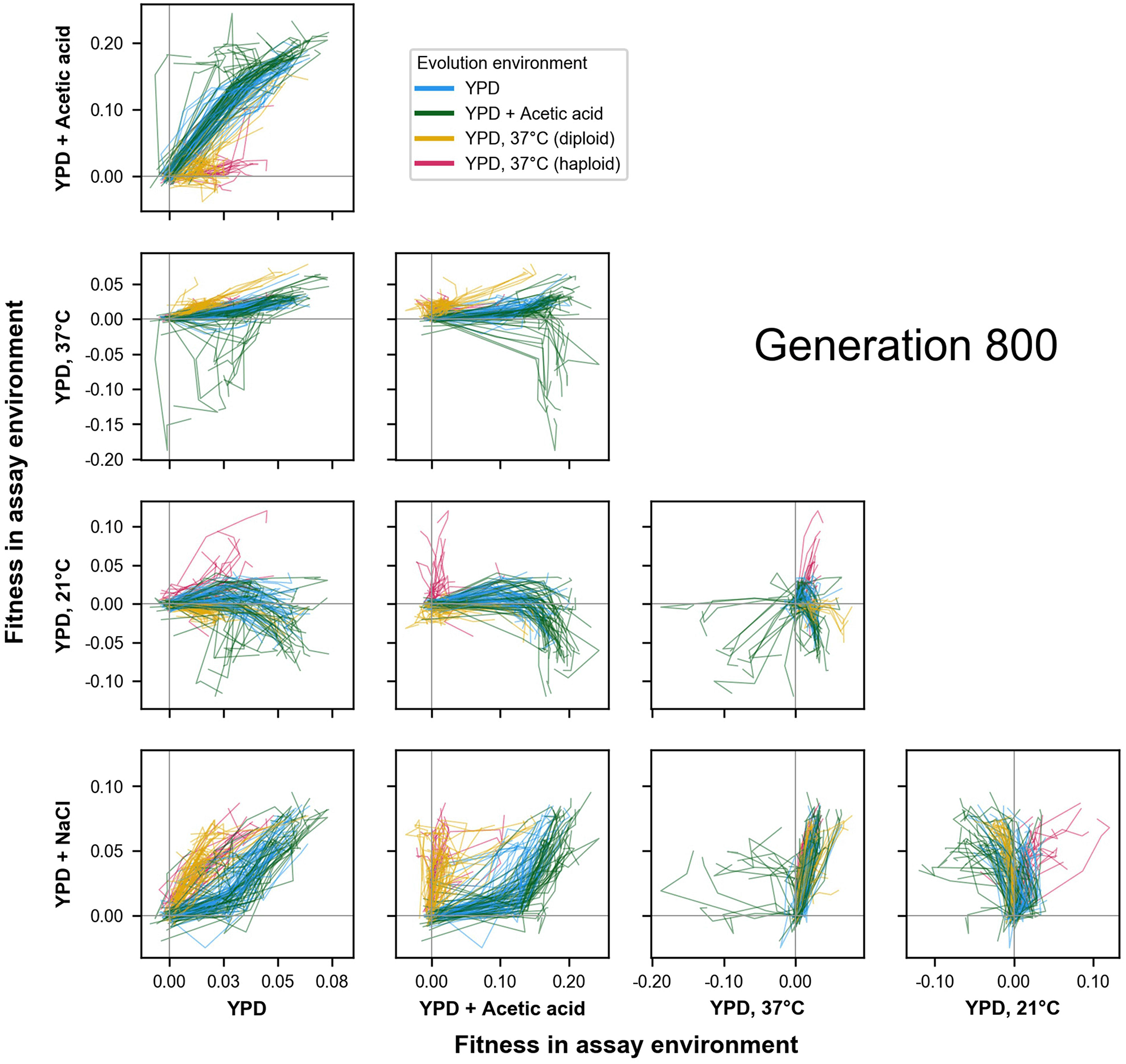

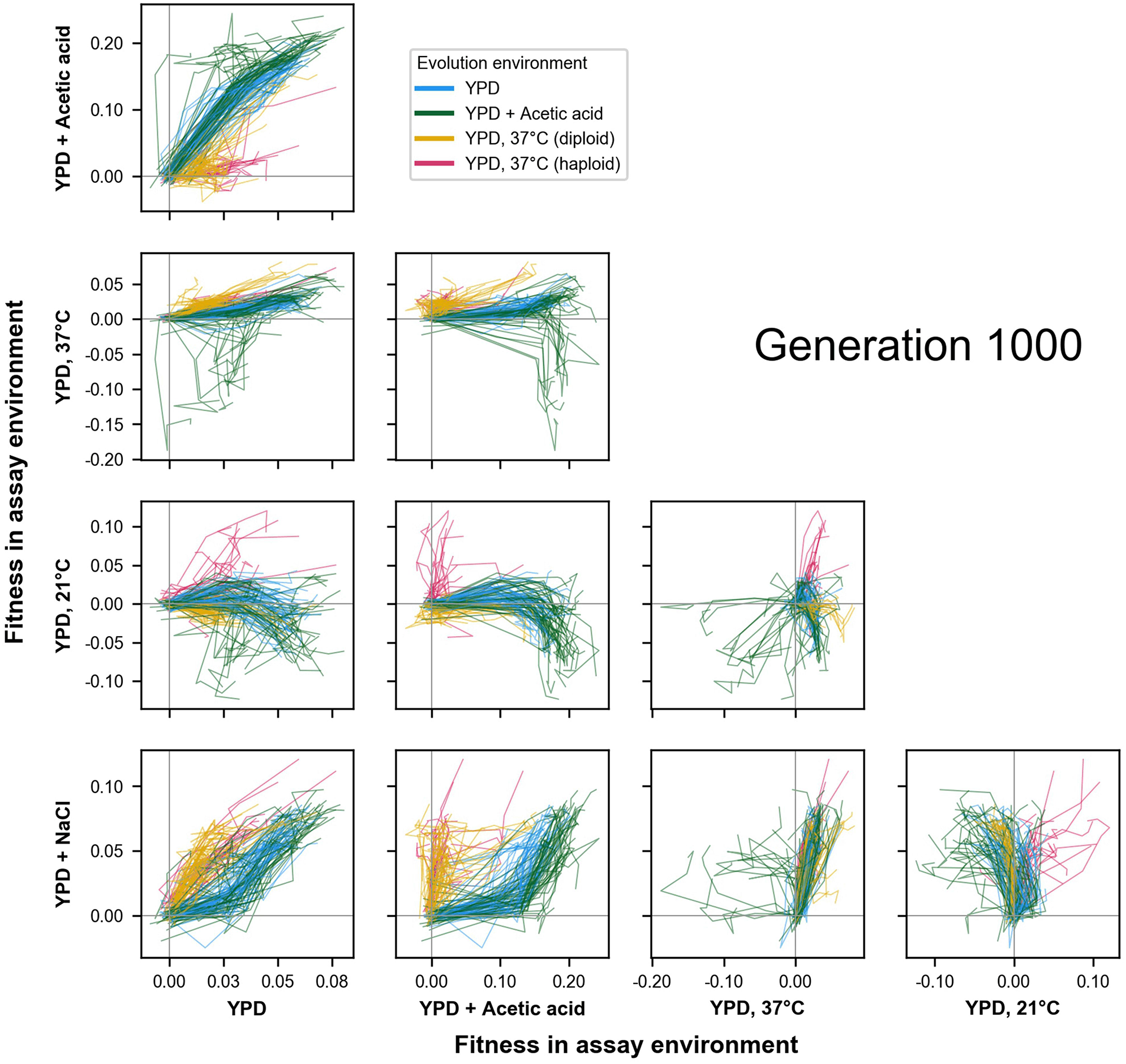

